# Comparative epigenomics across the barley pangenome links structural variation to regulatory genome function

**DOI:** 10.64898/2026.07.24.740502

**Authors:** Zihao Zhu, Erwang Chen, Pavla Navratilova, Miriam Schreiber, Sudharsan Padmarasu, Patrick König, Axel Himmelbach, Malcolm Macaulay, Robbie Waugh, Martin Mascher, Nils Stein

## Abstract

Structural variants (SVs) are abundant in plant genomes and influence agronomic traits, yet their regulatory interpretation remains challenging. Here, we combine pangenome-wide profiling of DNA methylation and chromatin accessibility across 20 barley genotypes, complemented by histone modification and chromatin interaction data in a subset of 10 genotypes. Comparative analysis of genotype-specific epigenomes reveals a globally conserved DNA methylation landscape across the barley pangenome alongside extensive regulatory variability at orthologous genes. We show that SVs do not broadly remodel global chromatin landscapes but instead act through context-dependent rewiring of local regulatory interactions. Despite this epigenomic stability, SVs may contribute to gene expression changes via chromatin contacts. Tissue-specific chromatin accessibility demonstrates that SV effects depend on developmental context. Integrating chromatin state variation with SVs at key vernalization genes explains epigenetic contributions to growth habit diversity. Together, these results provide a framework for interpreting the regulatory consequences of structural variation in crop genomes.

## Introduction

Structural variants (SVs), including presence-absence variation, copy number differences, and large chromosomal rearrangements, is a pervasive feature of eukaryotic genomes and a major driver of phenotypic diversity, environmental adaptation, and crop improvement^1^. Because single-reference genomes cannot capture the full extent of intraspecific variation, pangenomes that integrate multiple high-quality assemblies have emerged as essential resources for resolving genome diversity^2^. In barley (*Hordeum vulgare*), a diploid model for temperate cereals, successive pangenome efforts spanning 20 to 76 diverse wild and domesticated genotypes^3,4^, have uncovered extensive SVs associated with key agronomic traits, including disease resistance, plant architecture, seed dormancy^5^, and specialized metabolism^6^. Despite these advances, most pangenome studies have focused primarily on sequence variation, providing limited insight into the regulatory mechanisms through which SVs influence gene function. SVs have the potential to reshape gene regulation by altering the dosage, genomic context, and spatial organization of *cis*-regulatory elements, thereby reconfiguring chromatin architecture and long-range interactions^7^. These effects propagate across multiple regulatory layers including DNA methylation, chromatin accessibility, histone modifications, which together define chromatin states, as well as three-dimensional genome organization, which contributes to higher-order genome regulation^8^. In barley, single-reference epigenomic studies have revealed chromatin states with extensive distal interactions^9,10^. However, how these features vary across structurally diverse genomes remains unknown, underscoring the need to move beyond single-reference approaches to understand how SVs shape epigenomic landscapes within a pangenome. Here, we present a comparative epigenomics strategy that integrates pangenome-wide epigenomic profiling with comparative analyses across 20 barley genotypes, providing new insights into the regulatory consequences of SVs in crop genomes.

## Results

### Epigenomic profiling of twenty barley genomes

To systematically characterize regulatory features from DNA to higher-order chromatin organization and to extend beyond our previous single-reference datasets^10^, we applied sequencing-based approaches across the twenty genotypes of the barley pangenome version 1 (BPGv1)^4^, primarily using seedling leaf tissue. For all 20 genotypes, we profiled DNA methylation by whole-genome bisulfite sequencing (WGBS) and chromatin accessibility by assay for transposase-accessible chromatin using sequencing (ATAC-seq)^11^ **(Fig. 1a and Supplementary Table 1 and 2)**. In a subset of 10 genotypes, we further performed chromatin immunoprecipitation sequencing (ChIP-seq) for representative histone modifications (H3K4me3, H3K9ac, and H3K27me3)^12^, as well as promoter capture Hi-C **(Supplementary Table 3 and 4)**. To resolve three-dimensional (3D) genome organization, we generated Hi-C and/or Omni-C data in Morex, Barke, and RGT Planet, and additionally performed Hi-C and ATAC-seq in developing inflorescences of Morex and Barke. Given the extensive structural variation across the 20 genotypes, including large-scale inversions **(Extended Data Fig. 1)**, we constructed genotype-specific epigenomic profiles for each accession to minimize mapping bias.

**Fig. 1.**
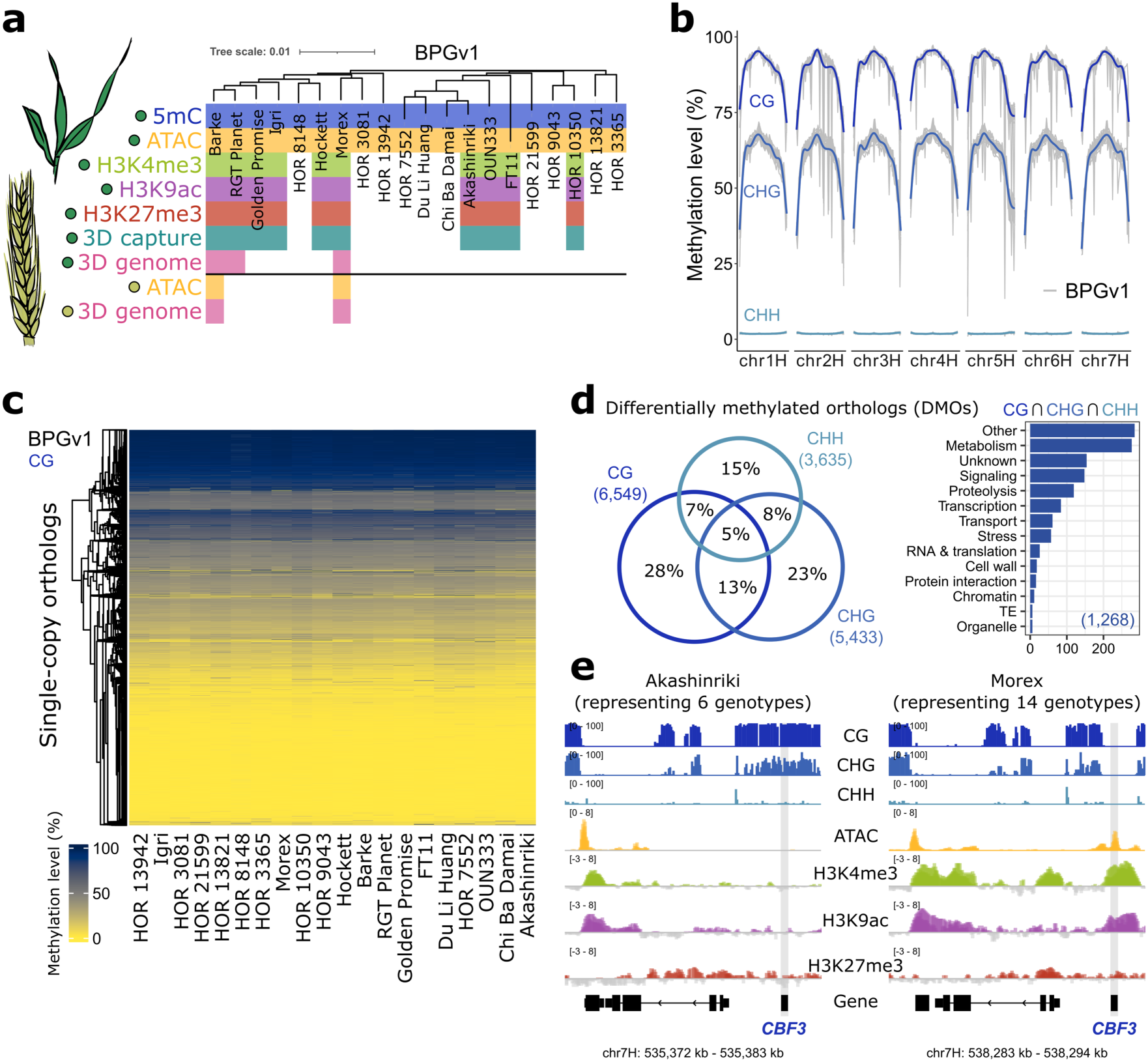
Epigenomic profiling and methylome variation across the barley pangenome. **a**, Overview of multi-layer epigenomic profiling across the barley pangenome, including DNA methylation, chromatin accessibility, histone modifications, and chromatin conformation assays in selected genotypes and tissues (seedling leaves and developing inflorescences). **b**, Chromosomal distribution of DNA methylation in CG, CHG, and CHH contexts across 20 barley genotypes (10 Mb bins); gray lines indicate individual genotypes and fitted curves show the overall trend. **c**, Heatmap of average CG methylation across 22,773 single-copy orthologs, clustered by methylation level. **d**, Overlap of differentially methylated orthologs (DMOs; Gaussian Mixture Model, ΔBIC > 10) across CG, CHG, and CHH contexts, together with Gene Ontology enrichment of shared DMOs **(Supplementary Data 2-4)**. **e**, Genome tracks of the *CBF3* locus showing genotype-specific CG and CHG methylation, chromatin accessibility, and histone modification signals; biological replicates are overlaid (ATAC-seq, n = 3; ChIP-seq, n = 2).

We first performed a comparative methylome analysis to investigate DNA methylation as a fundamental layer of epigenomic regulation^13^. While genotype-specific variation was evident, DNA methylation levels across all three sequence contexts (CG, CHG, and CHH) followed the expected chromosomal distribution patterns **(Fig. 1b)**. CG and CHG methylation were enriched in regions corresponding to putative centromeres and decreased toward chromosomal ends, whereas CHH methylation remained consistently low, consistent with patterns observed in other plant genomes^14,15^. Barley genomes are highly enriched in transposable elements (TEs), which comprise approximately 86% of the genome across all 20 genotypes **(Extended Data Fig. 2a and Supplementary Table 5)**. This high TE content is reflected in the elevated average methylation levels we observed in the CG (88.8%) and CHG (60.1%) contexts, which particularly for CG methylation, scale with genome size and TE content across species^16–20^ **(Extended Data Fig. 2b)**. TEs were preferentially methylated in CG and CHG contexts compared to gene bodies and coding sequences **(Extended Data Fig. 2c)**, in agreement with established patterns in plant epigenomes^21^. Together, these results indicate a conserved global methylation landscape across genotypes despite extensive structural variation.

Because our data were analyzed in genotype-specific coordinate systems, we integrated established orthologous relationships^3^ **(Supplementary Data 1)** to connect genes across genomes and assess context-specific variation in DNA methylation. Although average CG methylation levels across 22,773 single-copy orthologs were largely similar among genotypes **(Fig. 1c)**, 28.8% (6,549) of orthologs were differentially methylated in at least one genotype; we term these differentially methylated orthologs (DMOs). Among them, 2,896 were genotype-specific **(Extended Data Fig. 3a and Supplementary Data 2)**. Extending this analysis to CHG and CHH contexts identified 1,268 orthologs co-differentially methylated across all three contexts, affecting genes predominantly involved in core and secondary metabolic processes **(Fig. 1d and Extended Data Fig. 3b and Supplementary Data 3 and 4)**. Approximately 18% of total DMOs were shared between CG and CHG contexts, representing the largest pairwise overlap. One illustrative example is the barley homologue of the *Arabidopsis thaliana* cold acclimation-related *C-repeat Binding Factor 3* (*CBF3*) gene^22^, which is heavily methylated in CG and CHG contexts in the cultivar Akashinriki (and five other genotypes in the BPGv1 panel), but largely unmethylated in the remaining 14 genotypes, including Morex **(Fig. 1e and Supplementary Table 6)**. Similar DNA methylation variation was also observed at the related barley *CBF3-like* gene. In Akashinriki, methylation at the *CBF3* locus is accompanied by loss of chromatin accessibility and reduced levels of the active histone modification marks H3K4me3 and H3K9ac, consistent with coordinated epigenomic regulation^23^. To further investigate global relationships between DNA methylation and gene regulation, we correlated methylation levels with transcript abundance from the same tissue in the barley pan-transcriptome^24^ **(Extended Data Fig. 4a and Supplementary Data 5)**. This revealed a positive association for CG methylation, whereas CHG and CHH methylation showed the expected negative correlations, consistent with previous reports^21^ **(Extended Data Fig. 4b)**. Although the biological consequences for individual candidates (e.g., *CBF3*) require further investigation, these results demonstrate the utility of our comparative epigenomics for identifying candidate regulatory loci.

### Barley 3D genome organization and functional genome annotation

Having established locus level consistency, we next extended our analysis from gene-centric comparisons to genome-wide integration of epigenomic datasets. We investigated whether global DNA methylation levels are associated with chromatin accessibility, as previously observed in *Arabidopsis*^25^. Overlapping DNA methylation profiles with accessible chromatin regions (ACRs) identified by ATAC-seq revealed a clear inverse relationship, confirming that ACRs are largely depleted of DNA methylation^26^ **(Fig. 2a)**. As expected, most ACRs across all 20 genotypes were located in distal intergenic regions, promoters, 5’UTRs, and first exons. Both the genomic distribution of ACRs and their signal intensities, were highly conserved among genotypes **(Fig. 2b)**.

**Fig. 2.**
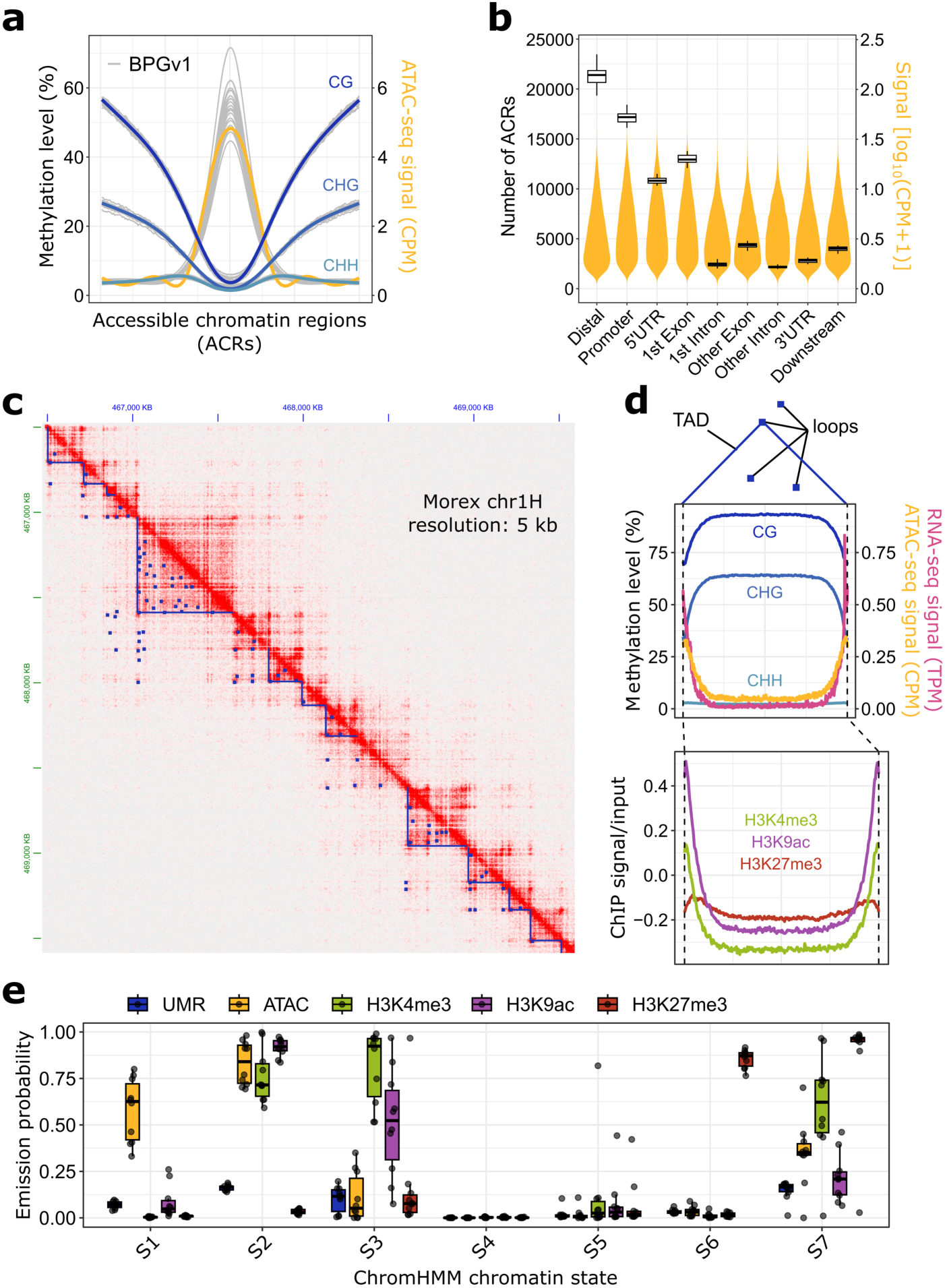
Integrated chromatin architecture and functional genome annotation in barley. **a**, Average DNA methylation and ATAC-seq signal profiles across accessible chromatin regions (ACRs; q < 0.05, ≥ 2 biological replicates); gray lines indicate individual genotypes (n = 20) and fitted curves show the overall trend. **b**, Distribution and signal intensity of ACRs across genomic features; boxplots show the ACR number across genotypes (n = 20) and violin plots display signal distributions of all detected ACRs with each genomic feature. **c**, Hi-C contact map of a representative region on Morex chr1H at 5 kb resolution, with annotated TADs (triangles) and chromatin loops (dots; q < 0.05). **d**, Average epigenomic and transcriptomic profiles of TADs in Morex (ATAC-seq and RNA-seq, n = 3; ChIP-seq, n = 2 biological replicates). **e**, ChromHMM chromatin state segmentation; boxplots show the distribution of scaled enrichment values (emission profiles) of unmethylated regions (UMRs), ATAC-seq signal, H3K4me3, H3K9ac, and H3K27me3 across the seven chromatin states (S1-S7) in 10 genotypes. Boxes represent medians and interquartile ranges, and whiskers extend to 1.5x the interquartile ranges.

Given that ACRs may possess *cis*-regulatory elements (CREs) involved in chromatin looping^27^, we next asked how these putative regulatory regions are organized in 3D nuclear space. A dense Hi-C map enabled the detection of chromatin loops and topologically associating domains (TADs) in the barley genome **(Fig. 2c)**, consistent with previous observations in hexaploid wheat^28,29^. In total 7,239 TADs with a median size of 190 kb were identified at 5 kb resolution, and were preferentially distributed along chromosome arms **(Extended Data Fig. 5a)**. TAD boundaries were characterized by reduced CG and CHG DNA methylation, elevated CHH methylation, increased chromatin accessibility and active transcription, and enrichment of histone modification, particularly H3K4me3 and H3K9ac **(Fig. 2d and Extended Data Fig. 5a)**; enrichment of H3K9ac at TAD boundaries has also been reported^28,30^. These features support that TAD formation is associated with transcriptionally active regions and may be driven, at least in part, by ACR-mediated *cis*-interactions^31^. Building on this observation, and recognizing that reliable TAD detection depends on sequencing depth **(Extended Data Fig. 5c and Supplementary Table 7)**, we used promoter-overlapping ACRs as anchors to generate capture Hi-C libraries. Capture Hi-C provides a cost-effective alternative to deeply sequenced Hi-C for interrogating chromatin interactions^32^. Although this approach does not achieve the same resolution for loop detection as deeply sequenced Hi-C maps, it retains the ability to detect key TAD structures **(Extended Data Fig. 5d)**.

Together, these results reveal the coordinated interplay among DNA methylation, chromatin accessibility and histone modifications within the context of 3D genome organization. To integrate these regulatory layers systematically, we combined all epigenomic datasets to define chromatin states and generate genome-wide functional annotations. Across the 10 genotypes with complete datasets, the genome was segmented into seven chromatin states using normalized enrichment profiles of five epigenomic features: unmethylated regions (UMRs), ATAC-seq signal, H3K4me3, H3K9ac, and H3K27me3 **(Extended Data Fig. 6a)**. Pairwise comparisons of chromatin state definitions showed highly similar enrichment patterns for each epigenomic feature across genotypes, indicating that the functional identity of individual states is largely preserved **(Extended Data Fig. 6b)**. State 1 was defined primarily by chromatin accessibility, with weak or limited enrichment of histone marks, consistent with unmodified ACRs^26^, whereas state 4 comprised highly methylated regions largely devoid of the assayed chromatin features. Conservation was most pronounced for the strong active chromatin state (state 2) and the Polycomb-repressed state (state 6), the latter marked by high H3K27me3 enrichment and low ATAC-seq signal, consistent with facultative chromatin **(Fig. 2e)**. In contrast, intermediate states (e.g., states 3 and 7) exhibited greater variation across genotypes, consistent with the possibility that these regulatory landscapes are more dynamic and susceptible to genotype- and tissue-specific reorganization.

### Epigenomic consequences of a large inversion

The pangenome-wide epigenomic datasets generated here enabled us to investigate the functional consequences of large structural variants (SVs), including a ∼141 Mb inversion on chromosome 7H (chr7H) identified in the cultivar RGT Planet relative to Morex and Barke^4^ **(Fig. 3a)**. Previous pan-transcriptome analyses revealed widespread transcriptional effects within this region and suggested that the inversion may influence 3D genome organization, as it spans boundary between an active (A) and inactive (B) compartment^24^. Comparative epigenomic profiling between RGT Planet and non-inverted genotypes (Morex and Barke) showed that both ACR distribution and DNA methylation patterns are reorganized in accordance with the inverted gene order, as well as the inverted distribution of Ty3 long terminal repeat (LTR) retrotransposons **(Extended Data Fig. 7a)**.

**Fig. 3.**
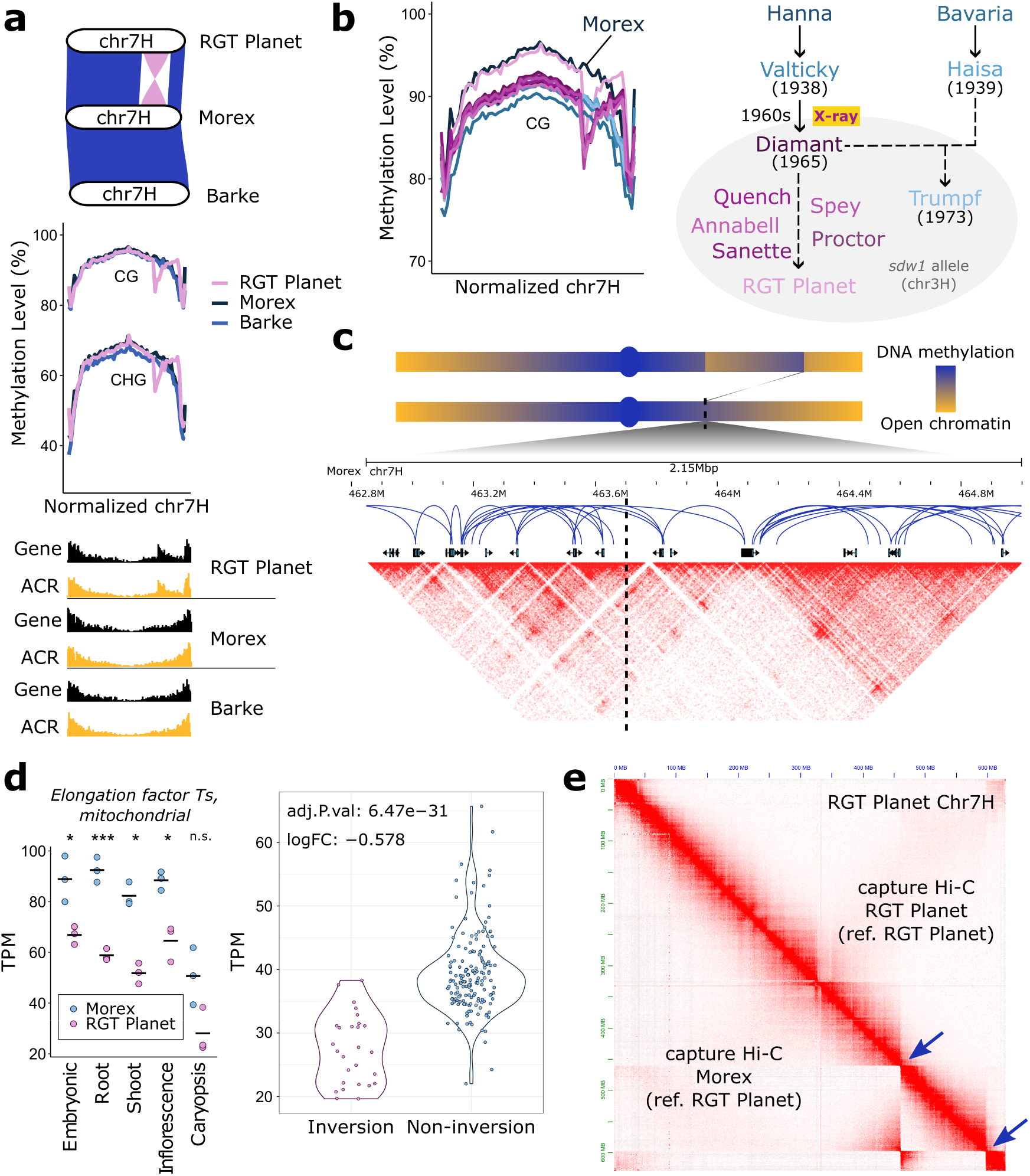
Epigenomic and chromatin interaction profiles across the chr7H inversion. **a**, Synteny plot of chr7H in Morex, Barke, and RGT Planet, ordered by gene position, with syntenic regions shown in blue and the inversion in pink; aligned tracks show DNA methylation (CG and CHG; 10 Mb bins), annotated genes, and ACRs across genotypes. **b**, Distribution of CG methylation on chr7H across barley cultivars, with pedigree relationships indicated (solid lines, direct parents; dashed lines, indirect parents). Genotypes shown in pink and purple colors carry the X-ray induced inversion, whereas those in blue do not. The gray region indicates genotypes carrying the recessive *sdw1* allele on chr3H. DNA methylation was profiled by WGBS for Morex and RGT Planet, and by PacBio 5mC detection for all other genotypes. **c**, Schematic model indicating that overall chromatin state remains stable despite the inversion. Hi-C contact map of Morex showing local chromatin interactions (annotated arcs, q < 0.05, 5 kb resolution) near the inversion breakpoint (dashed lines). **d**, TPM of the first gene downstream of the inversion breakpoint in Morex and RGT Planet ortholog across five tissues (n = 3 biological replicates, * *P* < 0.05, *** *P* < 0.001, n.s., not significant, two-tailed *t* test), and comparison of its TPM between genotypes with and without inversion from a European two-row spring barley population (n = 191). **e**, Capture Hi-C contact map of chr7H in RGT Planet (upper panel) and Morex (lower panel), both mapped to the RGT Planet reference genome; blue arrows indicate the inversion breakpoints.

Although RGT Planet is the only assembled genotype carrying this inversion in both BPGv1^4^ and BPGv2^3^, pedigree information traces its origin to X-ray-induced mutagenesis from Valticky to Diamant in the 1960s **(Fig. 3b)**. Despite strong selection for agronomic traits such as *sdw1*-mediated dwarfism^33^, recent extended pangenome analyses indicate that this inversion persists in multiple derived cultivars along the breeding lineage from Diamant to RGT Planet^34^. Notably, all genotypes carrying the inversion display highly similar CG methylation landscapes across the affected region, suggesting that global chromatin states remain largely stable across generations **(Fig. 3c)**, in agreement with observations from engineered inversions in *Arabidopsis*^35^. However, this genome-wide stability raises the possibility that higher-order chromatin structures such as TADs or chromatin loops are instead affected at finer resolution near inversion breakpoints. Indeed, the high-resolution Hi-C map in Morex revealed strong local interaction domains spanning this region, which are likely reorganized in RGT Planet due to the inversion **(Fig. 3c)**. Integrated analyses support this hypothesis: the gene closest to the breakpoint, annotated as a homologue of *Elongation factor Ts* in *Arabidopsis*^36^, shows only minor epigenomic changes **(Extended Data Fig. 7b)** but consistently reduced expression in RGT Planet across nearly all five tissues examined **(Fig. 3d)**. It is also classified as an inversion-responsive gene in a 191-cultivar spring barley panel **(Supplementary Data 6)**^24,37^.

Importantly, chromatin interactions involving breakpoint-proximal regions differ between capture Hi-C data from Morex mapped to the RGT Planet reference and capture Hi-C data from RGT Planet mapped to the same reference **(Fig. 3e)**. This reduction in interactions is confirmed by Omni-C data from RGT Planet **(Extended Data Fig. 7c)**, which indicates that the inversion region exhibits decreased contact frequency with adjacent chromosomal regions. Although plants lack canonical CTCF-mediated insulation, our results suggest that SVs can nonetheless alter local chromatin interaction landscapes, consistent with the broader principle that genome architecture can modulate gene regulation across eukaryotes^7^. These findings suggest that the chr7H inversion is associated with altered local chromatin interaction landscapes, which may contribute to transcriptional variation of genes in or near the affected region. However, effects on more distal genes are likely to involve additional regulatory mechanisms beyond local contact disruption. Because recombination is suppressed in the inverted region^4^, the inversion and its associated epigenomic features are inherited as a single linkage block, likely reflecting passive retention rather than direct selection, although selection on a linked secondary trait cannot be ruled out.

### Tissue-specific chromatin landscapes underlying gene regulation

The barley pan-transcriptome has revealed substantial tissue- and genotype-specific transcriptional complexity^24^. To assess whether this complexity is reflected, and potentially explained, at the epigenomic level, we performed comparative epigenomic analyses of Morex and Barke across two tissues and developmental stages, focusing on chromatin accessibility dynamics. We identified 40,419 and 38,424 genome-wide tissue-specific ACRs (TACRs; FDR < 0.05) between seedling leaves and developing inflorescences in Morex and Barke, respectively **(Fig. 4a and Extended Data Fig. 8a and Supplementary Data 7 and 8)**. These TACRs are distributed along chromosome arms, with a subset located within structural variation regions between Morex and Barke **(Fig. 4b)**.

**Fig. 4.**
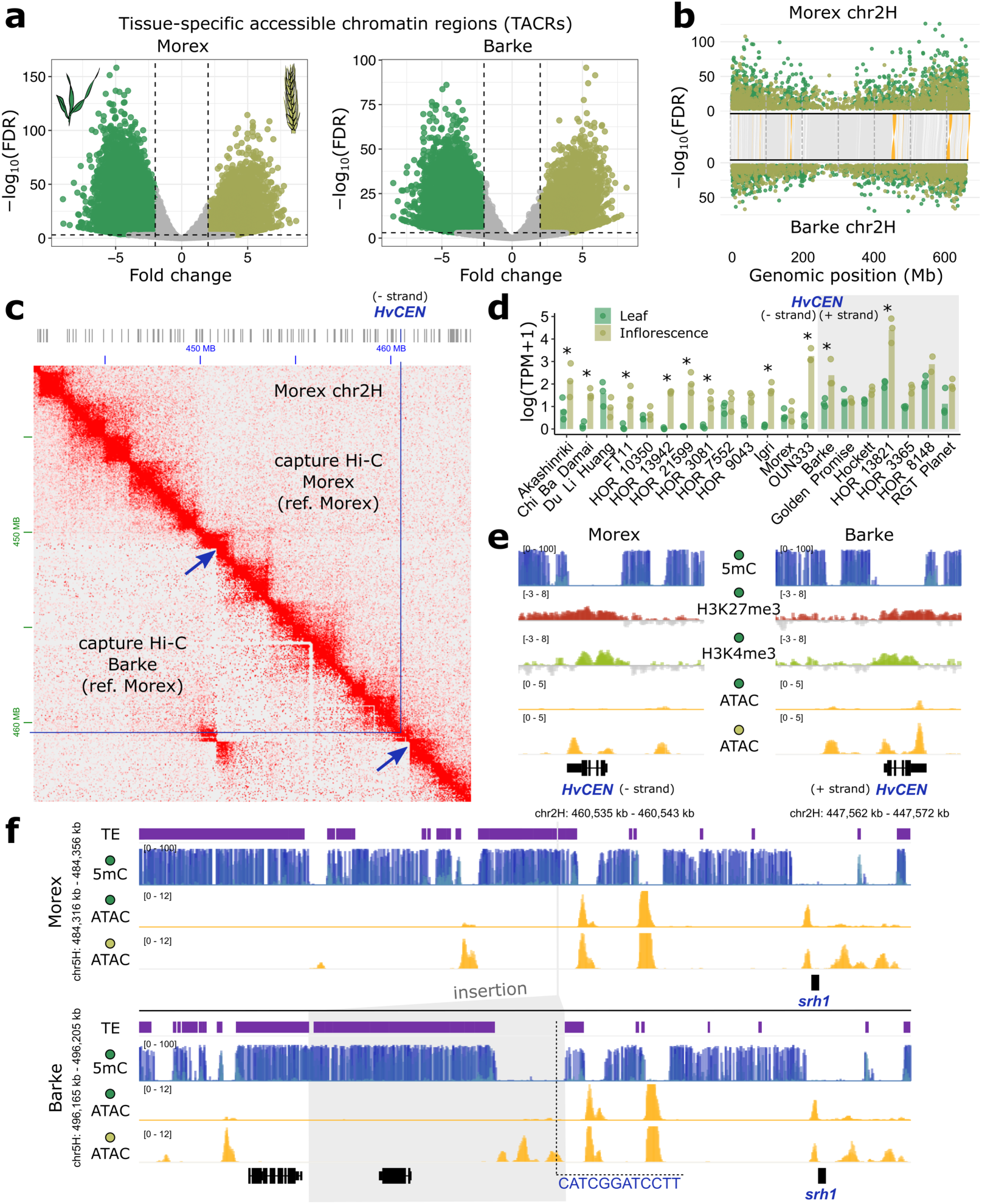
Tissue-specific chromatin accessibility and its association with transcription. **a**, Seedling leaf- and developing inflorescence-specific ACRs (FDR < 0.001, fold change > 2) in Morex and Barke. **b**, Distribution of tissue-specific ACRs (TACRs) across chr2H in Morex and Barke; inversions between Morex and Barke are shown in orange, with syntenic regions in gray. **c**, Capture Hi-C contact map spanning an inversion region in Morex (upper panel) and Barke (lower panel), including the *HvCEN* locus (blue lines) near the inversion breakpoint, both mapped to the Morex reference genome; blue arrows indicate the inversion breakpoints. **d**, TPM of *HvCEN* in seedling leaf and developing inflorescence across 20 barley genotypes. Bar charts indicate mean values (n = 2-3 biological replicates). Asterisks indicate significant differences between tissue types (*P* < 0.05, two-way ANOVA, **Supplementary Table 9**). The gray region indicates genotypes carrying the inversion (*HvCEN* on the negative strand). **e**-**f**, Genome tracks of the *HvCEN* (**e**) and *srh1* (**f**) loci in Morex and Barke showing DNA methylation (5mC, all contexts overlaid), histone modification signals (*HvCEN*, leaf), and chromatin accessibility (leaf and inflorescence); biological replicates are overlaid (ATAC-seq, n = 2-3; ChIP-seq, n = 2). **f**, The gray region indicates an insertion containing a putative enhancer in Barke.

One example is a ∼10 Mb inversion on chr2H, originally identified in domesticated barleys from northern Europe (e.g., in Barke), which has a breakpoint near the flowering-time repressor gene *CENTRORADIALIS* (*HvCEN*)^4,38^ **(Fig. 4c)**. Broader analysis across the 20 genotypes revealed that this inversion is not restricted to the original discovery set: seven genotypes carry the inverted Barke allele (+ strand), while 14 carry the non-inverted Morex allele (- strand) **(Fig. 4d and Supplementary Table 8)**. Although *HvCEN* expression is elevated in developing inflorescences overall, the inverted allele is generally associated with higher steady-state transcript levels^24^ (*P* < 0.001, **Supplementary Table 9**), consistent with its role as a flowering repressor and its contribution to delayed flowering. Correspondingly, the lower expression of the Morex allele is associated with reduced chromatin interactions and localization within a smaller TAD-like domain compared to the Barke allele in both tissue types **(Fig. 4c and Extended Data Fig. 8b)**. In seedling leaf tissue, the gene body and promoter of *HvCEN* are largely unmethylated; however, they exhibit concurrent enrichment of H3K4me3 and H3K27me3 together limited chromatin accessibility, corresponding to repressive and intermediate chromatin states **(Fig. 4e and Supplementary Table 8)**. By contrast, ATAC-seq data from developing inflorescences reveal strong accessibility at the gene termini and at a putative upstream regulatory element, indicating a more open chromatin state in both genotypes. This chromatin configuration is consistent with the observed tissue-specific expression patterns.

Beyond inversions, other SVs between Morex and Barke also contribute to tissue-specific regulatory variation, including an insertion originally identified in Barke that is associated with the long rachilla hair phenotype in the barley pangenome accessions^3^. Integration of comparative epigenomics data showed that this insertion region harbors a putative enhancer, is largely devoid of TEs, and remains unmethylated **(Fig. 4f)**. Notably, ACRs overlapping this enhancer are detected specifically in Barke developing inflorescences, but not in seedling leaf tissue. This tissue-specific accessibility mirrors the expression pattern of the target *srh1* gene^3^, consistent with the regulation of its *Arabidopsis* homologue *SIAMESE-RELATED* (*SMR*) by a similar enhancer motif^39,40^, and is consistent with a model in which dynamic enhancer accessibility contributes to the regulation of rachilla hair development in barley. Together, these results demonstrate that structural variation and tissue-specific chromatin dynamics jointly define regulatory landscapes across the barley pangenome.

### Epigenomic basis for growth habit variation

Barley exhibits substantial diversity in winter and spring growth habit, with the requirement for a cold period before flowering (vernalization) linked to the epigenetic regulation of the key vernalization gene *VRN1* conserved in wheat and barley^41–43^. Although the spring growth habit has traditionally been attributed to deletions of the flowering repressor *VRN2*, recent barley pangenome analyses show that *VRN2* is retained in many spring varieties, potentially reflecting inconsistencies in growth habit classification^44^. To refine growth habit classification across the pangenome, we performed flowering time experiments on 20 genotypes with and without standard vernalization under controlled conditions **(Extended Data Fig. 9a)**. This enabled a more nuanced classification based on vernalization responsiveness, whereby a subset of previously defined winter or spring barleys, including HOR 21599, Akashinriki, Chi Ba Damai, and HOR 13942, was reclassified as facultative. These accessions flower without cold exposure but still respond to vernalization (mean difference > 10 days; **Fig. 5a and Supplementary Table 9**), highlighting the importance of facultative types in buffering increasingly variable and unpredictable environmental conditions^45^. Facultative types also tend to exhibit increased vegetative growth in the absence of vernalization, as reflected by tiller number at heading **(Extended Data Fig. 9a and Supplementary Table 9)**. Consistent with this refined classification, spring types exhibit detectable steady-state *VRN1* transcript levels in seedling leaves prior to vernalization **(Fig. 5b)**. While *VRN1* expression distinguishes spring from winter and facultative barleys, it shows considerable variation among spring genotypes.

**Fig. 5.**
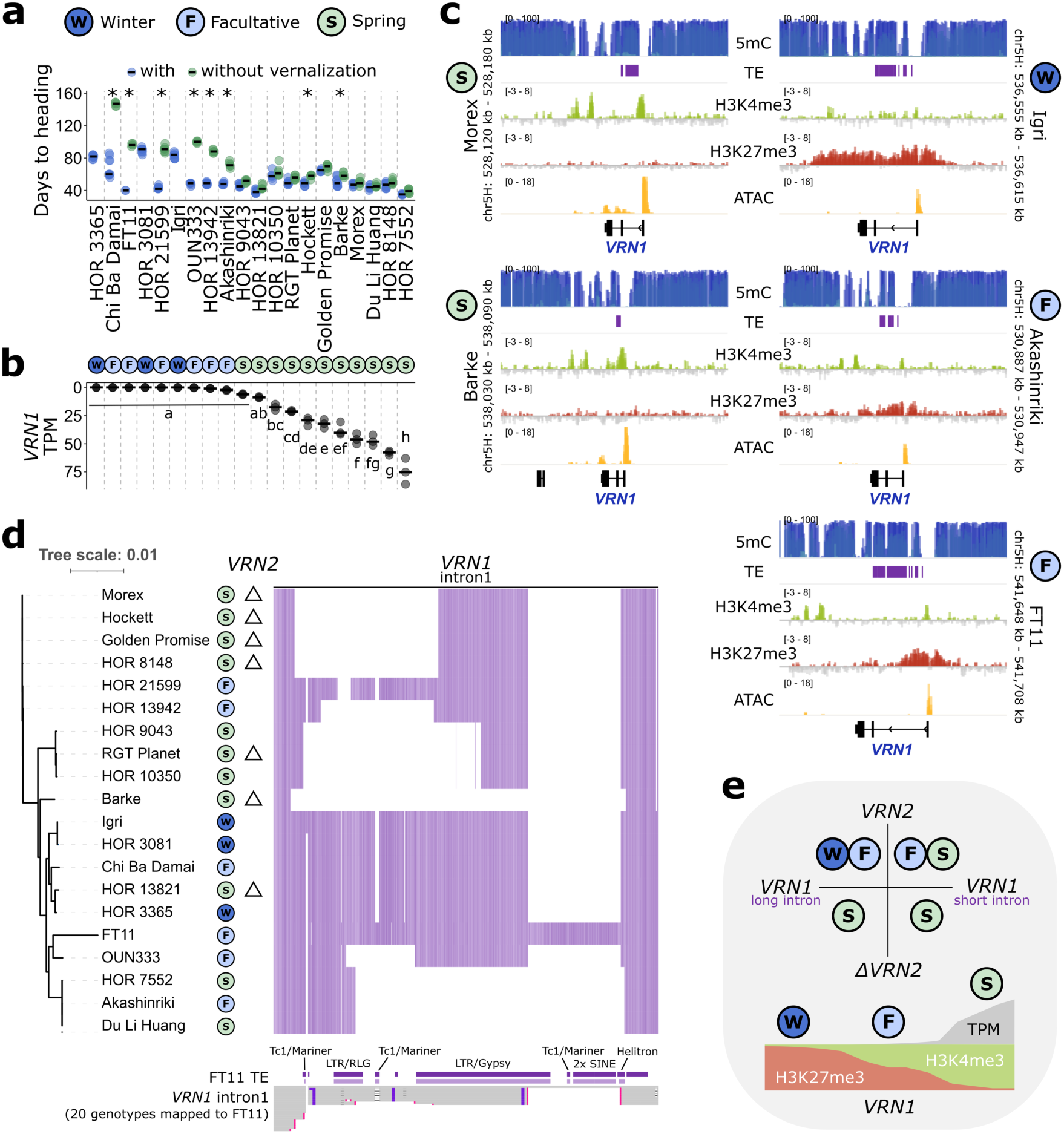
Epigenomic basis of vernalization requirements across the barley pangenome. **a**, Days to heading of plants with and without a six-week vernalization treatment. Horizontal bars indicate mean values (n = 2-10 biological replicates). Plants that did not head by day 160 without vernalization are classified as winter (W); plants showing a vernalization response (mean difference > 10; * *P* < 0.05, two-way ANOVA) but still heading without vernalization are classified as facultative (F); plants that do not respond to vernalization (mean difference ≤ 10) are classified as spring (S) **(Supplementary Table 9)**. **b**, TPM of *VRN1* in seedling leaf without vernalization treatment (n = 2-3 biological replicates). Different letters indicate significant differences (*P* < 0.05, one-way ANOVA followed by Tukey’s HSD test). **c**, Genome tracks of the *VRN1* locus in selected genotypes showing DNA methylation (5mC, all contexts overlaid), intron 1 TE annotation, histone modification signals, and chromatin accessibility; biological replicates are overlaid (ATAC-seq, n = 3; ChIP-seq, n = 2-3). **d**, Phylogenetic tree of *VRN1* (including 4 kb upstream region) across 20 barley genotypes, annotated with *VRN2* deletion status and *VRN1* intron 1 sequence alignment, including FT11 TE annotation (dark purple, raw annotation; light purple, curated annotation). The mapping track indicates TE presence/absence across genotypes. **e**, Proposed models linking growth habit classification in barley to genomic, transcriptomic, and epigenomic features.

Epigenomic profiling revealed distinct chromatin marks at the *VRN1* locus between winter and spring types. The winter barley cultivar Igri is enriched for the repressive histone mark H3K27me3, whereas the spring cultivars Morex and Barke show enrichment of the active mark H3K4me3 prior to vernalization **(Fig. 5c and Extended Data Fig. 9b)**. The chromatin state in winter barleys is consistent with previous reports in barley and wheat^41,42^, whereas the active chromatin configuration in spring barleys, consistent with increased chromatin accessibility and *VRN1* expression, suggests that these genotypes bypass vernalization-induced epigenetic reprogramming. Notably, facultative types (e.g., FT11 and Akashinriki) exhibit a mixed chromatin state, with both marks present at lower levels and more restricted regions **(Fig. 5c and Extended Data Fig. 9c)**. This pattern resembles the bivalent chromatin states of the key vernalization gene *FLOWERING LOCUS C* (*FLC*) in *Arabidopsis*^46^, although *FLC* and *VRN1* respond to vernalization in opposite directions, with *FLC* being repressed and *VRN1* activated. Together, these results confirm chromatin state at the *VRN1* locus as a key epigenetic determinant of growth habit variation across the barley pangenome.

Given the clear intron length variation observed across these genotypes, we systematically examined *VRN1* sequence variation across 76 BPGv2 assemblies^3^. Consistent with previous studies^47^, the primary source of variation resides in the first intron, a region prone to size expansion or contraction **(Fig. 5d and Extended Data Fig. 10 and Supplementary Table 10)**. For example, the wild barley (*Hordeum vulgare* subsp. *spontaneum*) accession FT11 contains an intron exceeding 14.5 kb, compared to the shortest intron approximately 1.8 kb in Barke. Repeat annotation indicates that this variation is largely driven by TE insertions. Notably, insertions located near the 5’ end of intron 1, including an LTR/Gypsy element, overlap H3K27me3-enriched regions in FT11, suggesting that TE-derived sequences represent candidate *cis*-elements that may facilitate the recruitment of repressive chromatin marks at *VRN1* and contributing to the previously observed but unresolved length variation at this locus. Importantly, long intron 1 alleles are mainly observed in winter and facultative barleys, with the spring barley accession HOR 13821 being a notable exception. Since this genotype carries a deletion of *VRN2*, these observations are broadly consistent with a prominent role of *VRN2* in determining growth habit in barley. Additionally, *VRN1* chromatin and sequence variation appear to represent an ancillary layer that might offer predictive value for vernalization requirements at the seedling stage **(Fig. 5e)**.

## Discussion

Several cereal crop species now possess reference pangenomes that capture extensive SVs across diverse germplasm, in some cases linking these variants to pan-transcriptomic differences in gene expression^24,48^. In parallel, pan-transcriptomic studies have revealed pervasive tissue- and genotype-specific transcriptional complexity. However, as these datasets primarily represent downstream transcriptional outputs, they offer limited insight into the regulatory mechanisms connecting genome structure to gene activity. Here, we integrate barley pangenome and pan-transcriptome resources within a multi-layered epigenomic framework, moving beyond descriptive catalogues of structural variation toward a unified view of regulatory genome organization. Our results show that SVs do not broadly remodel chromatin landscapes at the genome-wide scale but instead are associated with highly context-dependent regulatory effects, supporting emerging models from mammalian systems in which gene regulation is governed by the spatial organization of regulatory elements rather than linear genome structure^7^. In this framework, SVs can act as architectural modifiers that rewire regulatory interactions while largely preserving global chromatin state.

At the local level, regulatory features are highly sensitive to genome rearrangements. In particular, inversions that reposition genes and disrupt chromatin contacts are associated with reproducible and directionally consistent changes in gene expression. These effects are strongly context dependent, as tissue-specific chromatin accessibility reveals candidate regulatory elements active only at specific developmental stages, explaining why the same SV can be associated with divergent transcriptional consequences across tissues. By integrating epigenomic profiles with pangenome and pan-transcriptome data, we provide a route to interpret complex traits as the outcome of multi-layered regulatory variation rather than single-level genomic changes.

Using this integrative framework, we establish an epigenomic classification of growth habit in barley. Growth habit variation is associated with distinct chromatin states at the *VRN1* locus, where transposon-associated intron expansion correlates with increased H3K27me3 deposition and reduced expression in winter and facultative types, indicating that structural and epigenetic variation jointly modulate vernalization requirements. Beyond these biological insights, the datasets generated in this study are made available to the community via PanBARLEX (https://panbarlex.ipk-gatersleben.de). This resource enables systematic exploration of regulatory variation, including previously uncharacterized differentially methylated regions and tissue-specific accessible chromatin regions, and supports data-driven dissection of complex traits by both researchers and breeders.

Although current sampling remains limited in developmental and environmental scope, and key chromatin marks, notably H3K9 methylation, are not yet incorporated, variability in epigenomic data quality across assays and genotypes complicates cross-genotype state alignment. Nevertheless, this work provides a foundation for a pan-epigenome framework in barley. Finally, as most conclusions remain correlative, targeted genome engineering and perturbation experiments will be essential to establish causal links between structural variation, chromatin architecture, and gene expression, ultimately enabling predictive models for crop improvement.

## Methods

### Plant material and growth conditions

Twenty barley genotypes from the pangenome panel^4,24^ were obtained from the IPK Genebank, including Akashinriki, Barke, Chi Ba Damai, Du Li Huang, FT11, Golden Promise, Hockett, HOR 10350, HOR 13821, HOR 13942, HOR 21599, HOR 3081, HOR 3365, HOR 7552, HOR 8148, HOR 9043, Igri, Morex, OUN333, and RGT Planet.

For seedling leaf sampling, plants were grown in 35-well trays in controlled growth chambers under a 16 h light / 8 h dark photoperiod at 20°C (day) and 16°C (night) for two weeks. For developing inflorescences, plants were grown in a greenhouse under the same photoperiod and temperature regime. Inflorescences were collected at Waddington stages 6.0-7.0 (approximately 1.5 cm length, excluding awns). All samples were harvested 4 hours after lights-on, with two to five biological replicates collected depending on the experiment.

### Whole-genome bisulfite sequencing (WGBS)

Genomic DNA was extracted from 0.1 g leaf tissue using the DNeasy Plant Mini Kit (Qiagen, Germany). DNA was fragmented using a Covaris S220 focused ultrasonicator (Covaris, MA, USA) with settings of 160 W peak power, 8% duty factor, and 200 cycles for 60 s. Fragmented DNA was bisulfite-converted using the EZ DNA Methylation-Gold Kit (Zymo Research, CA, USA) and prepared for sequencing with the xGen Methyl-Seq DNA Library Prep Kit (IDT DNA, IA, USA), following the manufacturer’s instructions. Unmethylated Lambda DNA (Promega, WI, USA) was spiked in before bisulfite conversion to assess conversion efficiency. Libraries were paired-end sequenced (2 x 151 bp) on a NovaSeq 6000 system (Illumina, CA, USA) at IPK Gatersleben. Five biological replicates were included per genotype across all 20 genotypes **(Supplementary Table 1)**.

### Assay for transposase-accessible chromatin (ATAC-seq)

ATAC-seq was performed on fresh tissues immediately after harvesting, as described previously^10^. Briefly, 0.2 g of tissue was finely chopped with a razor blade in nuclei isolation buffer (0.25 M sucrose, 10 mM Tris-HCl, pH 8.0, 10 mM MgCl_2_, 1% Triton X-100, 5 mM β-mercaptoethanol) supplemented with 1x Halt™ Protease Inhibitor Cocktail (Thermo Fisher Scientific, MA, USA). The homogenate was filtered through a 50-μm cell strainer, and nuclei were washed twice with the same buffer before resuspension. For each extraction, an aliquot was analyzed by flow cytometry for quality control and quantification. Approximately 75,000 nuclei were aliquoted, and the nuclear pellet was resuspended in a transposition reaction containing Tagment DNA Enzyme TDE1 (Illumina, CA, USA) and incubated at 37°C for 30 min. Transposition DNA were purified using the MinElute PCR Purification Kit (Qiagen, Germany). Libraries were amplified using the NEBNext® High-Fidelity 2x PCR Master Mix (New England Biolabs, MA, USA), purified with VAHTS DNA Clean Beads (Vazyme, China), and paired-end sequenced (2 x 151 bp) on NovaSeq 6000 and NovaSeq X Plus systems (Illumina, CA, USA) at IPK Gatersleben. Three biological replicates were included for leaf samples across all 20 genotypes, and two biological replicates were included for developing inflorescences for Morex and Barke **(Supplementary Table 2)**.

### Chromatin immunoprecipitation (ChIP-seq) and CUT&Tag

The ChIP experiment was performed according to a previously described method^10^, with minor modifications, primarily involving the use of Universal Plant ChIP-seq Kit (Diagenode, Belgium) for reagents and consumables. Briefly, 3 g of fresh leaves were crosslinked for 15 min under vacuum in 1% formaldehyde (Sigma-Aldrich, MO, USA). Crosslinking was quenched by a 5-min incubation in 0.125 M glycine. The crosslinked tissues were pulverized in liquid nitrogen for nuclei extraction. The nuclear pellet was resuspended in sonication buffer, and chromatin was sheared using a Covaris S220 focused ultrasonicator (Covaris, MA, USA) with settings of 175 W peak power, 20% duty factor, and 200 cycles for 250 s. Immunoprecipitation was performed following the manufacturer’s instructions using the respective antibodies: anti-H3K4me3 (ab213224, Abcam, UK; Mp 04-745, Merck Millipore, Germany; C15410003, Diagenode, Belgium), anti-H3K9ac (ab4441, Abcam, UK), and anti-H3K27me3 (ab6002, Abcam, UK; C15410195, Diagenode, Belgium). ChIP-seq libraries, including input samples, were prepared using the NEBNext® Ultra™ II DNA Library Prep Kit for Illumina (New England Biolabs, MA, USA) and paired-end sequenced (2 x 151 bp) on NovaSeq 6000 and NovaSeq X Plus systems (Illumina, CA, USA) at IPK Gatersleben. Two to three biological replicates were included per antibody per genotype across 10 genotypes. Details of antibody usage per sample are provided in **Supplementary Table 3**.

For Morex and Barke leaf samples, one additional CUT&Tag library per antibody was prepared using the CUTANA™ CUT&Tag Kit (EpiCypher, NC, USA) following the manufacturer’s instructions. Nuclei extraction and quantification were performed as in the ATAC-seq experiment, and the following antibodies were used: anti-H3K4me3 (Mp 04-745, Merck Millipore, Germany), anti-H3K9ac (ab4441, Abcam, UK), and anti-H3K27me3 (13-0055t, EpiCypher, NC, USA). An anti-rabbit IgG antibody (13-0042t, EpiCypher, NC, USA) was used as a negative control. Libraries were paired-end sequenced (2 x 151 bp) on a NovaSeq 6000 system (Illumina, CA, USA) at IPK Gatersleben.

### Hi-C, capture Hi-C, and Omni-C sequencing

In situ Hi-C libraries were prepared from leaf samples of 10 genotypes, as well as from developing inflorescences of Morex and Barke, according to a previously described protocol using *Dpn*II for the digestion of crosslinked chromatin^49^. For capture Hi-C, in situ Hi-C libraries from leaf samples were enriched using the myBaits Hybridization Capture Kit (Daicel Arbor Biosciences, MI, USA) following the manufacturer’s instructions. Across the 10 genotypes, all ATAC-seq peaks overlapping 2-kb promoters were selected to generate 137,706 probes (each 120 bp in length) targeting accessible promoters. For Barke and RGT Planet leaf samples, additional Omni-C libraries were prepared using the Omni-C Kit (Dovetail Genomics, CA, USA) following the manufacturer’s instructions. Briefly, 0.5 g of snap-frozen samples was used as input, ground, and crosslinked with 3 mM DSG (Thermo Fisher Scientific, MA, USA), followed by 1% formaldehyde (Sigma-Aldrich, MO, USA). Chromatin was then digested using serial dilutions of the nuclease enzyme mix, and optimal digestion products were used for library preparation. Libraries were paired-end sequenced (2 x 151 bp or 2 x 111 bp) on NovaSeq 6000 system and NovaSeq X Plus systems (Illumina, CA, USA) at IPK Gatersleben and the James Hutton Institute (UK). Details of library types per genotype are provided in **Supplementary Table 4**.

### Primary sequencing data processing

Sequencing data were processed using established pipelines and accompanying scripts tailored to each sequencing assay. All analyses were performed using genotype-specific reference^3^, i.e., 20 genome assemblies corresponding to the 20 barley genotypes.

For WGBS, adapters were trimmed from raw sequencing files with Trim Galore (v0.6.4^50^ with the parameters “--illumina --paired --clip_R1 10 --clip_R2 10”. Trimmed files were mapped to the corresponding genome using the Bismark pipeline (v0.24.1)^51^. Five replicates per genotype were merged using SAMtools (v1.16.1)^52^, deduplicated, and unique reads with “-q 30” were kept for methylation calling with the parameters “--CX” to include all three methylation contexts. Genome-wide average methylation levels per genotype were calculated **(Supplementary Table 1)**. The coverage of methylation information per single base resolution per context was analyzed using MethylDackel (v0.6.1) (https://github.com/dpryan79/MethylDackel) and bedGraphToBigWig (v2.8)^53^.

For ATAC-seq, ChIP-seq, and CUT&Tag, reads were adapter-trimmed using fastp (v0.20.0)^54^, mapped to the corresponding genome using BWA-MEM (v0.7.17)^55^, deduplicated, and quality filtered with “-q 30” using SAMtools (v1.16.1)^52^. Blacklisted regions were identified per genotype to exclude regions with abnormally high coverage from downstream analyses. For ChIP-seq, 5-kb bins with Z-score >=5 in input samples were defined as blacklisted regions. For ATAC-seq, 5-kb bins within the top 0.05% of coverage (>= 99.95th percentile) were defined as blacklisted regions. Peak calling was performed with “-q 0.05” using MACS (v3.0.0)^56^. For ChIP-seq, peaks were called against the corresponding input samples to identify narrow peaks for H3K4me3 and H3K9ac and broad peaks for H3K27me3 with “--broad-cutoff 0.1”. Coverage tracks were generated using deepTools (v3.5.1)^57^: bamCoverage with counts per million (CPM) normalization for ATAC-seq and CUT&Tag, and bamCompare for ChIP-seq by comparing IP and input samples **(Supplementary Table 2 and Supplementary Table 3)**.

For Hi-C, capture Hi-C, and Omni-C, reads were adapter-trimmed using fastp (v0.20.0)^54^. Mapping and filtering were performed using the HiC-Pro (v3.1.0) pipeline^58^. Briefly, reads were assigned to ligation fragments based on DpnII restriction sites (Hi-C and capture Hi-C) or the bridge sequence (Omni-C). Invalid pairs, including self-ligated fragments, unligated fragments, and duplicates were removed, and valid interaction pairs from biological replicates were merged for downstream analyses **(Supplementary Table 4)**.

### Identification of differentially methylated orthologs (DMOs)

All DNA methylation analyses were conducted in a context-specific manner. Genome-wide distributions of DNA methylation levels (10-Mbp window), as well as methylation levels across gene and transposable element (TE) regions, were calculated using ViewBS (version 0.1.9)^59^ and visualized using custom R scripts. TEs were de novo annotated across 20 genotypes based on genome assemblies and gene annotations using the EDTA (v2.2.2) pipeline^60^ **(Supplementary Table 5)**. Average methylation levels per gene were calculated based on the intersection of methylation coverage and gene annotations using BEDtools (version 2.30.0)^61^. The average transcript abundance per gene^24^ was integrated with methylation levels based on the gene annotation. Across the 20 genotypes, methylation information was integrated using established single-copy orthologous groups^3^ **(Supplementary Data 1)** and visualized using the R package ComplexHeatmap^62^. Orthologs with more than two genotypes missing methylation values were excluded from downstream analyses. To identify differentially methylated genes across orthologs and genotypes, Gaussian mixture models (GMMs) with one and two components were fitted independently to each ortholog based on average methylation levels, and model selection was based on the difference in Bayesian Information Criterion (ΔBIC). Orthologs showing strong support for two components (ΔBIC > 10) were classified as DMOs, while all others were assigned to a single cluster representing similarly methylated orthologs **(Supplementary Data 2-4)**. Cluster assignments from the two GMM-derived groups were converted into a binary matrix, with the smaller cluster labeled as 1 and the larger cluster (up to 19 genotypes) labeled as 0, and were sorted for visualization using the R package pheatmap^63^. Epigenomic profiles of the identified DMOs and their flanking regions were visually inspected using IGV^64^.

### ChromHMM chromatin state analysis

To functionally annotate each genome by integrating leaf epigenomic datasets, ChromHMM (v1.23)^65^ was applied following a previously described pipeline for Morex^10,66^. Briefly, unmethylated regions (UMRs) were identified by calculating average methylation levels in 100-bp bins using BEDtools (version 2.30.0)^61^, followed by filtering bins with <= 1% methylation and removing isolated or non-contiguous bins. Adjacent low-methylation bins were then merged, regions shorter than 300 bp were excluded and the remaining were extended to 200 bp minimum to define final UMRs. Together with BAM files from ATAC-seq and ChIP-seq, UMRs were binarized into 200-bp bins to learn seven chromatin sates, followed by overlap enrichment analysis. To assess the fold enrichment of each state across genomic features, annotation categories, including distal regions, promoters, 5’UTRs, first exons, first introns, other exons, other introns, 3’UTRs, and downstream regions, were generated based on gene annotation^3^.

### Identification of topologically associating domains (TADs) and chromatin loops

Valid interaction pairs generated by HiC-Pro (v3.1.0) were converted for visualization using hicpro2juicebox.sh^58^. To assess the effect of sequencing depth on TAD and loop detection, in silico subsampling was performed on the Morex leaf Hi-C dataset (approximately 25.9 billion valid interaction pairs) **(Supplementary Table 7)**. For deeply sequenced Morex and Barke leaf samples, three-dimensional genome features were identified using HOMER (v.4.10) findTADsAndLoops.pl^67^ with the parameters “-res 5000 -window 10000 -maxDist 1000000” and “-balance” for Hi-C matrix normalization. For capture Hi-C and developing inflorescence samples, the parameters were adjusted to “-res 10000 -window 20000” and “-res 20000 -window 50000” to account for differences in data resolution. Loop calling was performed using p-value threshold of 0.05. Detected TADs and loops were annotated and validated with the contact map in Juicebox (v2.17.00) with “balance++” normalization^68^. TAD boundary features from the high-resolution Morex leaf dataset were characterized by integrating newly generated epigenomic data with previously published RNA-seq datasets^69^ (PRJEB14349) using deepTools (v3.5.1) computeMatrix^57^.

### Breakpoint verification of inversions on chr2H and chr7H

The impact of chromosomal inversions on gene positions across 20 genotypes ^4^ was assessed by constructing a synteny network of orthologous genes using GENESPACE^70^ in combination with OrthoFinder (v2.5.5)^71^. To precisely define inversion breakpoints, whole-genome alignments were performed between Morex and Barke for the chr2H inversion, and between Morex and RGT Planet for the chr7H inversion, using minimap2 (v2.24)^72^. Genomic coordinates of the inversion breakpoints were subsequently identified with SyRI (v1.6)^73^.

### Analysis of the large inversion on chr7H

Eleven breeding founder lines related to RGT Planet were analyzed, including Annabell, Bavaria, Diamant, Haisa, Hanna, Proctor, Quench, Sanette, Spey, Trumpf, and Valticky. Information on the chromosome 7H inversion was derived from genome assemblies^34^. PacBio Hifi reads were mapped to the corresponding genomes using ccsmeth (v0.5.0)^74^ with default parameters. Methylation calling was performed using pb-CpG-tools (v6.4.0) (https://github.com/PacificBiosciences/pb-CpG-tools). Genome-wide CG methylation levels (10-Mbp window) were calculated using BEDtools (version 2.30.0)^61^ and visualized using custom R scripts.

Transcript variation of genes related to the chr7H inversion was analyzed by comparing Morex and RGT Planet pan-transcriptome datasets^24^ across five tissues. In addition, the crown RNA-seq data from the BARN dataset^37^ (PRJEB49069) was analyzed using a RGT Planet-specific reference transcript dataset (RTD)^24^. This BARN dataset includes 191 cultivated two-row European spring barley genotypes, of which 27 carry the chr7H inversion. Transcript abundance was estimated using Salmon^75^, and lowly expressed genes were filtered out (CPM > 2 in at least 25 genotypes), resulting in 20,602 expressed genes, including 845 in the inversion region. Differential expression between inversion and non-inversion genotypes was analyzed in R using the edgeR and limma-voom pipeline^76^, filtering for an adjusted p-value of ≤ 0.001 and a log fold change > 0.5, identified 318 differentially expressed genes (117 within the inversion) **(Supplementary Data 6)**.

### Identification of tissue-specific accessible chromatin regions (TACRs)

From Morex and Barke leaf and developing inflorescence samples, significant ATAC-seq peaks presenting ACRs, as well as corresponding BAM files, were used as input to identify TACRs per genotype using the R packages DiffBind^77^ and DESeq2^78^, with an FDR threshold of 0.05 **(Supplementary Data 7 and 8)**. TACRs with FDR < 0.001 and fold change > 2 were selected for allele mining and chromosomal distribution analysis. Epigenomic profiles of the identified TACRs and their flanking regions were visually inspected using IGV^64^, including regions containing the *HvCEN* locus^4^ and the *srh1* locus^3^.

### Growth habit classification

Growth habit of 20 genotypes was classified based on flowering time with and without vernalization treatment. Ten seeds per genotype were grown in 35-well trays in a greenhouse under controlled conditions with a 16 h light / 8 h dark photoperiod at 16°C (day) and 14°C (night) for two weeks, before being transferred to a standard vernalization treatment at constant 4°C with a 10 h light / 14 h dark photoperiod for six weeks. The non-vernalized batch was sown six weeks later to ensure that all plants were at similar developmental stages when transferred to 11 L pots and placed under control conditions. Non-germinated plants were excluded from further analyses, and all plants were randomly rotated twice per week. Days to flowering (first visible awns) were scored on a daily basis, and total tiller number per plant at flowering was counted. For the vernalized batch, the vernalization period was excluded from the calculation of days to flowering. Plants that did not show visible awns within 160 days were considered as non-flowering.

### Analysis of vernalization genes

The presence or absence of *VRN2* and steady-state transcript levels of *VRN1* in seedling shoot tissues (pre-vernalization) across 20 genotypes were obtained from previous analyses^44^ and published datasets^24^. Across 76 barley genomes^3^, *VRN1* variation was assessed using a similar approach **(Supplementary Table 10)**. Briefly, *VRN1* alleles were identified through sequence homology searches using BLASTn (v2.13.0)^79^, with AY758233.1^80^ as the reference sequence. Sequence alignments were performed using MUSCLE^81^ in AliView^82^, and visualized using the R package ggmsa^83^. Phylogenetic trees were constructed from full-length *VRN1* genomic sequences from 76 genotypes, and from *VRN1* genomic sequences including 4 kb upstream regions from 20 genotypes, using IQ-Tree (v2.2.2.6)^84^ under the HKY+F+G4 model, with 1,000 ultrafast bootstrap replicates. *VRN1* intron 1 sequences were further analyzed for TE insertions using separate EDTA (v.2.2.2)^60^ runs with a curated library TREP (https://trep-db.uzh.ch), classifying TEs into distinct categories. Epigenomic profiles of *VRN1* and its flanking regions were visually inspected using IGV^64^.

## Data availability

All raw data are available through the ENA (https://www.ebi.ac.uk/ena/browser/home), including WGBS reads (PRJEB95102), ATAC-seq reads (PRJEB98256), ChIP-seq/CUT&Tag reads (PRJEB106812), and chromatin conformation assay reads (PRJEB96274, PRJEB112837, and PRJEB111850). Visualizations of the processed epigenomic data for all genotypes at the gene level as well as in aggregated form at the level of orthologous groups (CDS clusters) are available at PanBARLEX (https://panbarlex.ipk-gatersleben.de).

## Code availability

Scripts used for data analysis are available at https://github.com/zihaozhu92/pepBAR.

## Author contributions

N.S. and M. Mascher conceived the project. Z.Z. and N.S. designed the study. Z.Z., S.P., M.S., and A.H. prepared epigenomic libraries and performed the sequencing. Z.Z. analyzed the sequencing data. E.C. analyzed transposable elements, structural variation, and methylomes in breeding cultivars. P.N. performed genome segmentation analysis. M.S., M. Macaulay, and R.W. provided and analyzed BARN transcriptome data associated with the chr7H inversion. P.K. provided orthologous relationships and integrated the data into PanBARLEX. Z.Z. wrote the manuscript. All authors reviewed and revised the manuscript.

## Acknowledgements

We thank Pascal Jaroschinsky, Jörg Fuchs, Manuela Knauft, and Jacqueline Pohl for technical support, and Anne Fiebig for data submission. This work was supported by the German Federal Ministry of Research, Technology and Space (BMFTR) [grants Epic-p-epBAR (FKZ 031B1224), SHAPE (FKZ 031B0190A), SHAPE-P2 (FKZ 031B0884A), and SHAPE-P3 (FKZ 031B1302A)], the German Research Foundation (DFG) within the project “Establishment of the National Research Data Infrastructure (NFDI)” in the consortium DataPLANT (Project Number 442077441), and James Hutton Institute and the Scottish Government RESAS project KJHI-B1-2, BBSRC project (BB/X018636/1, BB/S020160/1, ERA-CAPS BB/S004610/1 and BB/S020160/1).

## Supplementary information

Supplementary Table 1. Summary statistics of WGBS

Supplementary Table 2. Summary statistics of ATAC-seq

Supplementary Table 3. Summary statistics of ChIP-seq and CUT&Tag

Supplementary Table 4. Summary statistics of capture Hi-C, Hi-C, and Omni-C

Supplementary Table 5. Genome-wide TE composition across 20 barley genotypes

Supplementary Table 6. Differentially methylated *CBF3* and *CBF3L* across 20 barley genotypes

Supplementary Table 7. TAD and loop detection from chromatin conformation assays

Supplementary Table 8. *HvCEN* allelic orientation across 20 barley genotypes

Supplementary Table 9. Two-way ANOVA group means with Tukey’s HSD post-hoc comparisons

Supplementary Table 10. *VRN1* and *VRN2* variation across 76 barley genotypes

Supplementary Data 1. Single-copy orthologs across 20 barley genotypes

Supplementary Data 2. Differentially methylated orthologs in CG context

Supplementary Data 3. Differentially methylated orthologs in CHG context

Supplementary Data 4. Differentially methylated orthologs in CHH context

Supplementary Data 5. Average TPM of single-copy orthologs from seedling shoot samples

Supplementary Data 6. Differential expression associated with the chr7H inversion

Supplementary Data 7. Tissue-specific accessible chromatin regions in Morex

Supplementary Data 8. Tissue-specific accessible chromatin regions in Barke

## References

1 Zhu, Z., Jhingan, S., Chen, E. & Stein, N. Functional cereal pan-genomics: harnessing structural and regulatory variation for precision crop design. Current Opinion in Biotechnology 97, 103418 (2026).

2 Schreiber, M., Jayakodi, M., Stein, N. & Mascher, M. Plant pangenomes for crop improvement, biodiversity and evolution. Nature Reviews Genetics 25, 563–577 (2024).

3 Jayakodi, M. et al. Structural variation in the pangenome of wild and domesticated barley. Nature, 1–9 (2024).

4 Jayakodi, M. et al. The barley pan-genome reveals the hidden legacy of mutation breeding. Nature 588, 284–289 (2020).

5 Jørgensen, M. E. et al. Postdomestication selection of MKK3 shaped seed dormancy and end-use traits in barley. Science 391, 90–95 (2026).

6 Dias, S. L. et al. Biosynthesis of the allelopathic alkaloid gramine in barley by a cryptic oxidative rearrangement. Science 383, 1448–1454 (2024).

7 Spielmann, M., Lupiáñez, D. G. & Mundlos, S. Structural variation in the 3D genome. Nature Reviews Genetics 19, 453–467 (2018).

8 Domb, K., Wang, N., Hummel, G. & Liu, C. Spatial features and functional implications of plant 3D genome organization. Annual Review of Plant Biology 73, 173–200 (2022).

9 Baker, K. et al. Chromatin state analysis of the barley epigenome reveals a higher - order structure defined by H3K27me1 and H3K27me3 abundance. The Plant Journal 84, 111–124 (2015).

10 Navratilova, P. et al. Epigenome and interactome profiling uncovers principles of distal regulation in the barley genome. Cell Genomics 6 (2026).

11 Buenrostro, J. D., Wu, B., Chang, H. Y. & Greenleaf, W. J. ATAC-seq: a method for assaying chromatin accessibility genome-wide. Current protocols in molecular biology 109, 21.29. 21–21.29. 29 (2015).

12 Le, H., Simmons, C. H. & Zhong, X. Functions and mechanisms of histone modifications in plants. Annual Review of Plant Biology 76 (2025).

13 Xie, G., Du, X., Hu, H. & Du, J. Molecular mechanisms underlying the establishment, maintenance, and removal of DNA methylation in plants. Annual review of plant biology 76 (2025).

14 Cokus, S. J. et al. Shotgun bisulphite sequencing of the Arabidopsis genome reveals DNA methylation patterning. Nature 452, 215–219 (2008).

15 Lister, R. et al. Highly integrated single-base resolution maps of the epigenome in Arabidopsis. Cell 133, 523–536 (2008).

16 Feng, S. et al. Conservation and divergence of methylation patterning in plants and animals. Proceedings of the National Academy of Sciences 107, 8689–8694 (2010).

17 Gent, J. I. et al. CHH islands: de novo DNA methylation in near-gene chromatin regulation in maize. Genome research 23, 628–637 (2013).

18 Roelfs, K.-U., Känel, A., Twyman, R. M., Prüfer, D. & Schulze Gronover, C. Epigenetic variation in early and late flowering plants of the rubber-producing Russian dandelion Taraxacum koksaghyz provides insights into the regulation of flowering time. Scientific Reports 14, 4283 (2024).

19 Schmitz, R. J. et al. Epigenome-wide inheritance of cytosine methylation variants in a recombinant inbred population. Genome research 23, 1663–1674 (2013).

20 Stein, N., et al. Unveiling centromeric retrotransposon dynamics through a near-complete rye genome assembly. (2025).

21 Lloyd, J. P. & Lister, R. Epigenome plasticity in plants. Nature Reviews Genetics 23, 55–68 (2022).

22 Gilmour, S. J., Sebolt, A. M., Salazar, M. P., Everard, J. D. & Thomashow, M. F. Overexpression of the Arabidopsis CBF3 transcriptional activator mimics multiple biochemical changes associated with cold acclimation. Plant physiology 124, 1854–1865 (2000).

23 Xu, G. & Law, J. A. Loops, crosstalk, and compartmentalization: it takes many layers to regulate DNA methylation. Current opinion in genetics & development 84, 102147 (2024).

24 Guo, W. et al. A barley pan-transcriptome reveals layers of genotype-dependent transcriptional complexity. Nature Genetics, 1–10 (2025).

25 Zhong, Z. et al. DNA methylation-linked chromatin accessibility affects genomic architecture in Arabidopsis. Proceedings of the National Academy of Sciences 118, e2023347118 (2021).

26 Lu, Z. et al. The prevalence, evolution and chromatin signatures of plant regulatory elements. Nature plants 5, 1250–1259 (2019).

27 Ricci, W. A. et al. Widespread long-range cis-regulatory elements in the maize genome. Nature plants 5, 1237–1249 (2019).

28 Concia, L. et al. Wheat chromatin architecture is organized in genome territories and transcription factories. Genome Biology 21, 104 (2020).

29 Jia, J. et al. Homology-mediated inter-chromosomal interactions in hexaploid wheat lead to specific subgenome territories following polyploidization and introgression. Genome Biology 22, 26 (2021).

30 An, J. et al. An atlas of the tomato epigenome reveals that KRYPTONITE shapes TAD-like boundaries through the control of H3K9ac distribution. Proceedings of the National Academy of Sciences 121, e2400737121 (2024).

31 Liu, C., Cheng, Y.-J., Wang, J.-W. & Weigel, D. Prominent topologically associated domains differentiate global chromatin packing in rice from Arabidopsis. Nature plants 3, 742–748 (2017).

32 Šimková, H., Câmara, A. S. & Mascher, M. Hi-C techniques: from genome assemblies to transcription regulation. Journal of Experimental Botany 75, 5357–5365 (2024).

33 Dockter, C. & Hansson, M. Improving barley culm robustness for secured crop yield in a changing climate. Journal of experimental botany 66, 3499–3509 (2015).

34 Marone, M. P. et al. Selecting genomes that matter: haplotype-based prioritization for iterative pangenome expansion. bioRxiv, 2026.2005. 2013.724976 (2026).

35 Khosravi, S. et al. Epigenetic state and gene expression remain stable after CRISPR/Cas-mediated chromosomal inversions. New Phytologist 245, 2527–2539 (2025).

36 Browning, K. S. The plant translational apparatus. Plant molecular biology 32, 107–144 (1996).

37 Schreiber, M. et al. Genomic resources for a historical collection of cultivated two-row European spring barley genotypes. Scientific Data 11, 66 (2024).

38 Comadran, J. et al. Natural variation in a homolog of Antirrhinum CENTRORADIALIS contributed to spring growth habit and environmental adaptation in cultivated barley. Nature genetics 44, 1388–1392 (2012).

39 Kumar, N. et al. Functional conservation in the SIAMESE-RELATED family of cyclin-dependent kinase inhibitors in land plants. The Plant Cell 27, 3065–3080 (2015).

40 Nomoto, Y. et al. A hierarchical transcriptional network activates specific CDK inhibitors that regulate G2 to control cell size and number in Arabidopsis. Nature Communications 13, 1660 (2022).

41 Liu, Y. et al. Epigenomic identification of vernalization cis-regulatory elements in winter wheat. Genome Biology 25, 200 (2024).

42 Oliver, S. N., Finnegan, E. J., Dennis, E. S., Peacock, W. J. & Trevaskis, B. Vernalization-induced flowering in cereals is associated with changes in histone methylation at the VERNALIZATION1 gene. Proceedings of the National Academy of Sciences 106, 8386–8391 (2009).

43 Zeng, R. et al. Temperature regulation in plants: From molecular mechanisms to climate-resilient crop improvement. Journal of Integrative Plant Biology (2026).

44 Zhu, Z. & Stein, N. Pangenome insights into structural variation and functional diversification of barley CCT motif genes. The Plant Genome 18, e70098 (2025).

45 Hirsz, D. et al. VERNALIZATION2 alters early tiller development in a facultative spring hexaploid bread wheat. New Phytologist (2026).

46 Gao, Z. et al. A pair of readers of bivalent chromatin mediate formation of Polycomb-based “memory of cold” in plants. Molecular Cell 83, 1109–1124. e1104 (2023).

47 Hemming, M. N., Fieg, S., James Peacock, W., Dennis, E. S. & Trevaskis, B. Regions associated with repression of the barley (Hordeum vulgare) VERNALIZATION1 gene are not required for cold induction. Molecular Genetics and Genomics 282, 107–117 (2009).

48 Avni, R. et al. A pangenome and pantranscriptome of hexaploid oat. Nature, 1–9 (2025).

49 Padmarasu, S., Himmelbach, A., Mascher, M. & Stein, N. In situ Hi-C for plants: an improved method to detect long-range chromatin interactions. Methods Mol. Biol 1933, 441–472 (2019).

50 Krueger, F. Trim Galore https://github.com/FelixKrueger. TrimGalore> accessed 8 (2023).

51 Krueger, F. & Andrews, S. R. Bismark: a flexible aligner and methylation caller for Bisulfite-Seq applications. bioinformatics 27, 1571–1572 (2011).

52 Li, H. et al. The sequence alignment/map format and SAMtools. bioinformatics 25, 2078–2079 (2009).

53 Kent, W. J., Zweig, A. S., Barber, G., Hinrichs, A. S. & Karolchik, D. BigWig and BigBed: enabling browsing of large distributed datasets. Bioinformatics 26, 2204–2207 (2010).

54 Chen, S., Zhou, Y., Chen, Y. & Gu, J. fastp: an ultra-fast all-in-one FASTQ preprocessor. Bioinformatics 34, i884–i890 (2018).

55 Vasimuddin, M., Misra, S., Li, H. & Aluru, S. in 2019 IEEE international parallel and distributed processing symposium (IPDPS). 314–324 (IEEE).

56 Zhang, Y. et al. Model-based analysis of ChIP-Seq (MACS). Genome biology 9, R137 (2008).

57 Ramírez, F. et al. deepTools2: a next generation web server for deep-sequencing data analysis. Nucleic acids research 44, W160 (2016).

58 Servant, N. et al. HiC-Pro: an optimized and flexible pipeline for Hi-C data processing. Genome biology 16, 259 (2015).

59 Huang, X., Zhang, S., Li, K., Thimmapuram, J. & Xie, S. ViewBS: a powerful toolkit for visualization of high-throughput bisulfite sequencing data. Bioinformatics 34, 708–709 (2018).

60 Ou, S. et al. Benchmarking transposable element annotation methods for creation of a streamlined, comprehensive pipeline. Genome biology 20, 275 (2019).

61 Quinlan, A. R. & Hall, I. M. BEDTools: a flexible suite of utilities for comparing genomic features. Bioinformatics 26, 841–842 (2010).

62 Gu, Z. Complex heatmap visualization. Imeta 1, e43 (2022).

63 Kolde, R. & Kolde, M. R. Package ‘pheatmap’. R package 1, 790 (2015).

64 Robinson, J. T. et al. Integrative genomics viewer. Nature biotechnology 29, 24–26 (2011).

65 Ernst, J. & Kellis, M. Chromatin-state discovery and genome annotation with ChromHMM. Nature protocols 12, 2478–2492 (2017).

66 Crisp, P. A. et al. Stable unmethylated DNA demarcates expressed genes and their cis-regulatory space in plant genomes. Proceedings of the National Academy of Sciences 117, 23991–24000 (2020).

67 Heinz, S. et al. Simple combinations of lineage-determining transcription factors prime cis-regulatory elements required for macrophage and B cell identities. Molecular cell 38, 576–589 (2010).

68 Durand, N. C. et al. Juicebox provides a visualization system for Hi-C contact maps with unlimited zoom. Cell systems 3, 99–101 (2016).

69 Mascher, M. et al. A chromosome conformation capture ordered sequence of the barley genome. Nature 544, 427–433 (2017).

70 Lovell, J. T. et al. GENESPACE tracks regions of interest and gene copy number variation across multiple genomes. elife 11, e78526 (2022).

71 Emms, D. M. & Kelly, S. OrthoFinder: solving fundamental biases in whole genome comparisons dramatically improves orthogroup inference accuracy. Genome biology 16, 157 (2015).

72 Li, H. Minimap2: pairwise alignment for nucleotide sequences. Bioinformatics 34, 3094–3100 (2018).

73 Goel, M., Sun, H., Jiao, W.-B. & Schneeberger, K. SyRI: finding genomic rearrangements and local sequence differences from whole-genome assemblies. Genome biology 20, 277 (2019).

74 Ni, P. et al. DNA 5-methylcytosine detection and methylation phasing using PacBio circular consensus sequencing. Nature communications 14, 4054 (2023).

75 Patro, R., Duggal, G., Love, M. I., Irizarry, R. A. & Kingsford, C. Salmon provides fast and bias-aware quantification of transcript expression. Nature methods 14, 417–419 (2017).

76 Law, C. W. et al. RNA-seq analysis is easy as 1-2-3 with limma, Glimma and edgeR. F1000Research 5, ISCB Comm J-1408 (2018).

77 Stark, R. & Brown, G. DiffBind: differential binding analysis of ChIP-Seq peak data. R package version 100, 2–21 (2011).

78 Love, M., Anders, S. & Huber, W. Differential analysis of count data–the DESeq2 package. Genome Biol 15, 10–1186 (2014).

79 Camacho, C. et al. BLAST+: architecture and applications. BMC bioinformatics 10, 1–9 (2009).

80 von Zitzewitz, J. et al. Molecular and structural characterization of barley vernalization genes. Plant molecular biology 59, 449–467 (2005).

81 Edgar, R. C. MUSCLE: multiple sequence alignment with high accuracy and high throughput. Nucleic acids research 32, 1792–1797 (2004).

82 Larsson, A. AliView: a fast and lightweight alignment viewer and editor for large datasets. Bioinformatics 30, 3276–3278 (2014).

83 Zhou, L. et al. ggmsa: A visual exploration tool for multiple sequence alignment and associated data. Briefings in Bioinformatics 23, bbac222 (2022).

84 Minh, B. Q. et al. IQ-TREE 2: new models and efficient methods for phylogenetic inference in the genomic era. Molecular biology and evolution 37, 1530–1534 (2020).

