## Extended Data Figures 1-10 for "Comparative epigenomics across the barley pangenome links structural variation to regulatory genome function"

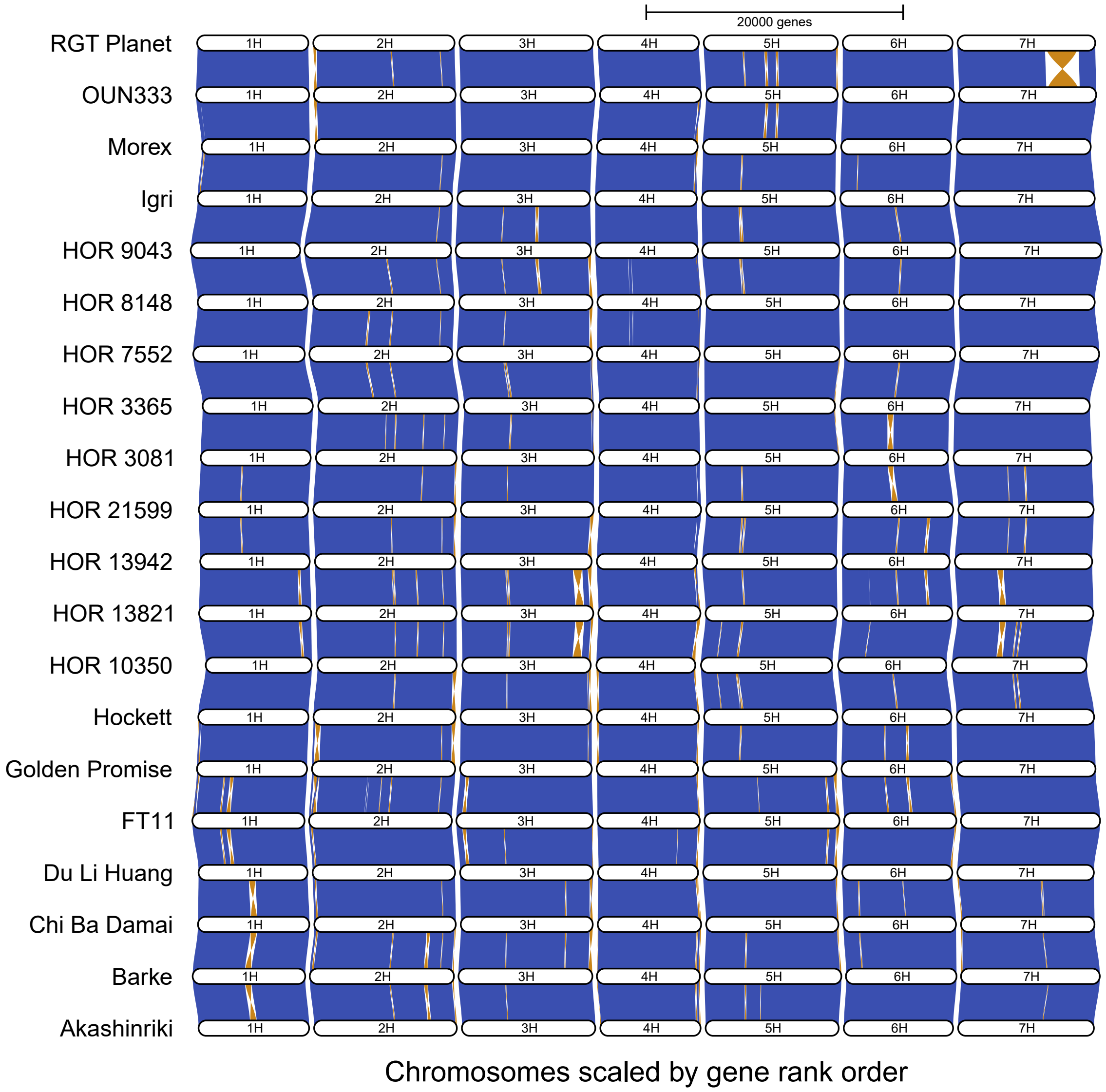


**Extended Data Fig. 1. Inversions affecting gene order across 20 genotypes.**
GENESPACE synteny plot showing chr1H to chr7H across 20 barley genotypes, ordered by gene position. Syntenic regions are shown in blue, and inversions are highlighted in orange.

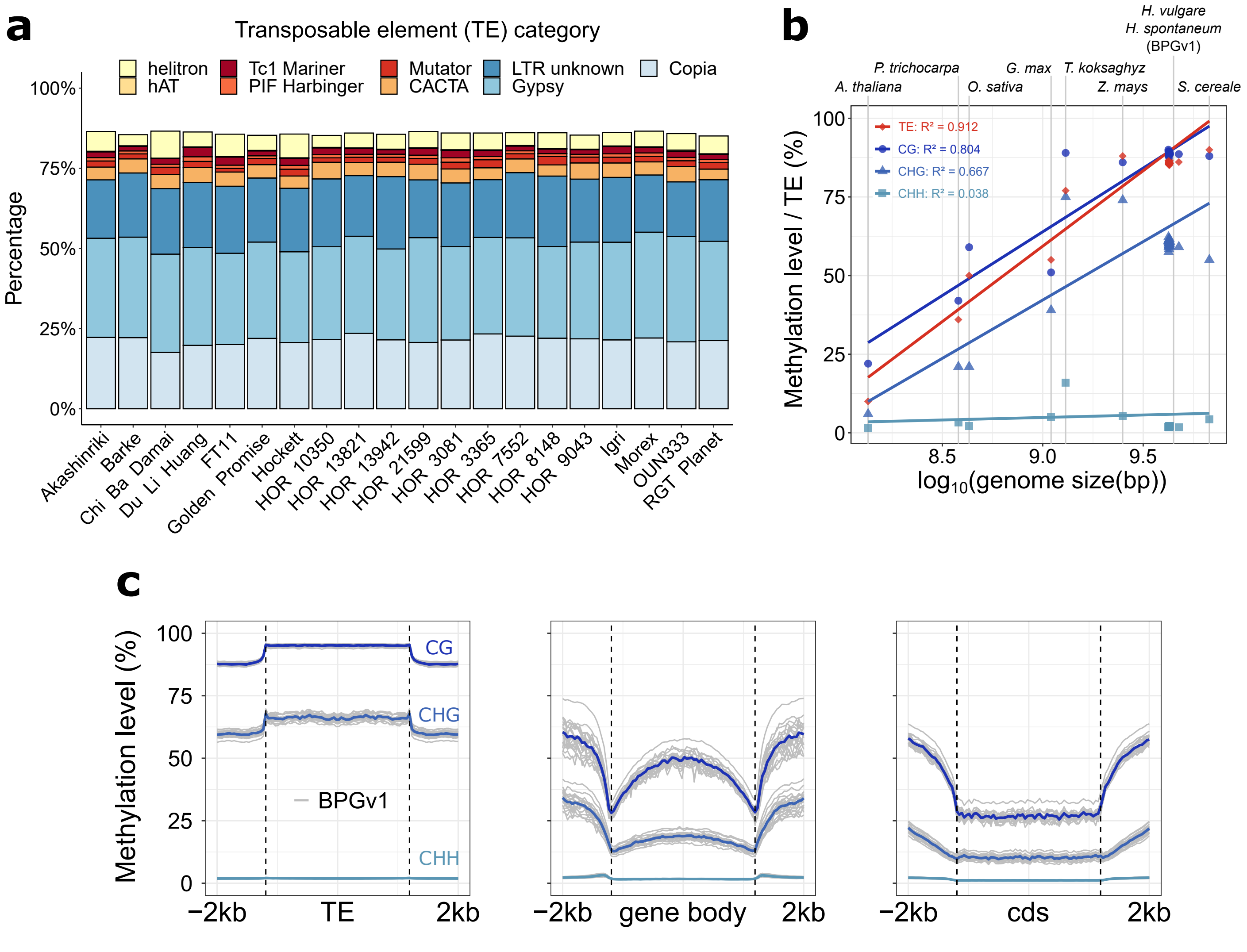


**Extended Data Fig. 2. Association of DNA methylation levels and transposable elements.**
**a**, De novo transposable element (TE) annotation across 20 barley genotypes **(Supplementary Table 5)**. **b**, Correlations between TE proportion, average DNA methylation levels (CG, CHG, and CHH contexts), and genome size across selected plant species. From left to right: *Arabidopsis thaliana*, *Populus trichocarpa*, *Oryza sativa*, *Glycine max*, *Taraxacum koksaghyz*, *Zea mays*, BPGv1 (including 19 *Hordeum vulgare* and 1 *Hordeum spontaneum*; average values across all BPGv1 accessions were used for correlation analyses; *H. vulgare* genotype HOR 8148 has a slightly larger genome size), and *Secale cereale*. **c**, Average DNA methylation profiles across TEs, gene bodies, and coding sequences; gray lines indicate individual genotypes (n = 20) and fitted curves show the overall trend.

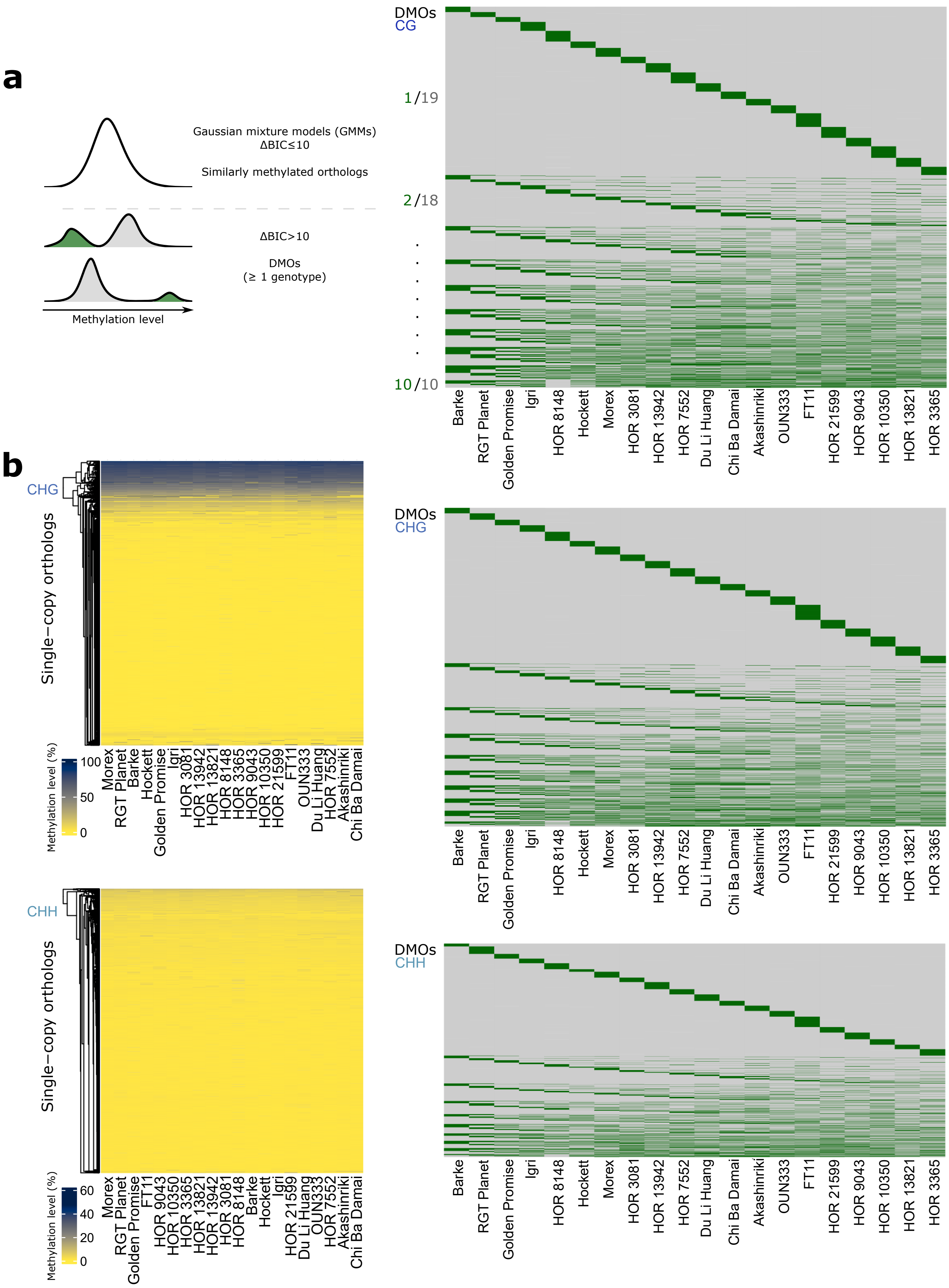


**Extended Data Fig. 3. Differentially methylated orthologs across 20 genotypes.**
**a**, Differentially methylated orthologs (DMOs) were identified using a Gaussian Mixture Model with two clusters (ΔBIC > 10) based on their average methylation levels. DMO clustering map (CG methylation) is shown based on the number of genotypes clustered in the same group (e.g., 1/19 indicates genotype-specific DMOs). **b-c**, Heatmaps of average CHG (**b**) and CHH (**c**) methylation across single-copy orthologs, clustered by methylation level, with corresponding DMO clustering maps. Clustering of DMOs for all contexts is provided in **Supplementary Data 2-4**.

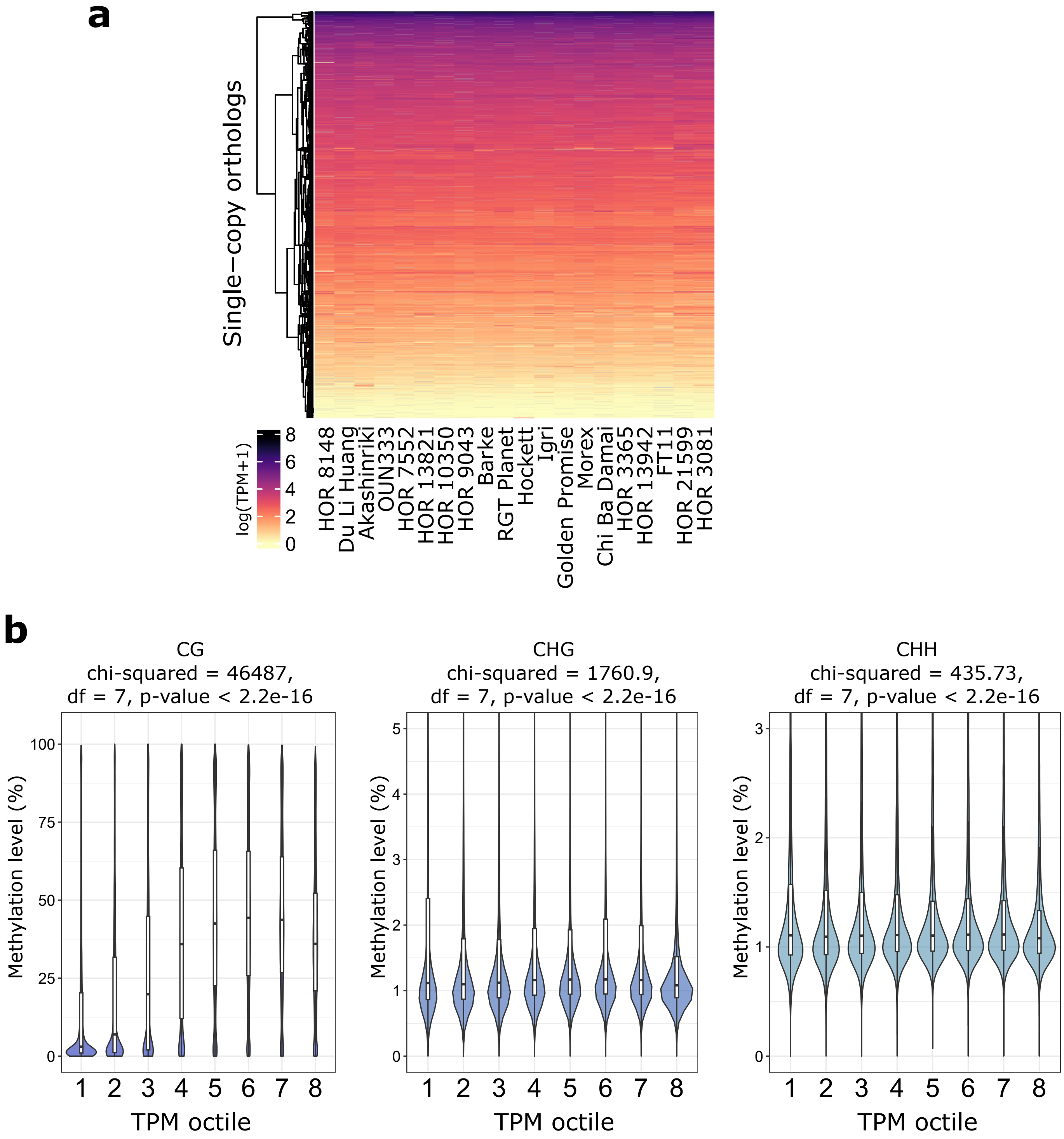


**Extended Data Fig. 4. Association of DNA methylation and transcript abundance.**
**a**, Heatmap of transformed average TPM across 10,135 single-copy orthologs, clustered by transformed average TPM value **(Supplementary Data 5)**. **b**, Distribution of average methylation levels (CG, CHG, and CHH contexts) across TPM octile (1-8, from low to high abundance). Violin and boxplots show all genes with available TPM data across 20 genotypes (n = 270,011). Boxes represent medians and interquartile ranges, and whiskers extend to 1.5x the interquartile ranges. Significant differences in distribution patterns were assessed using the Kruskal-Wallis test.

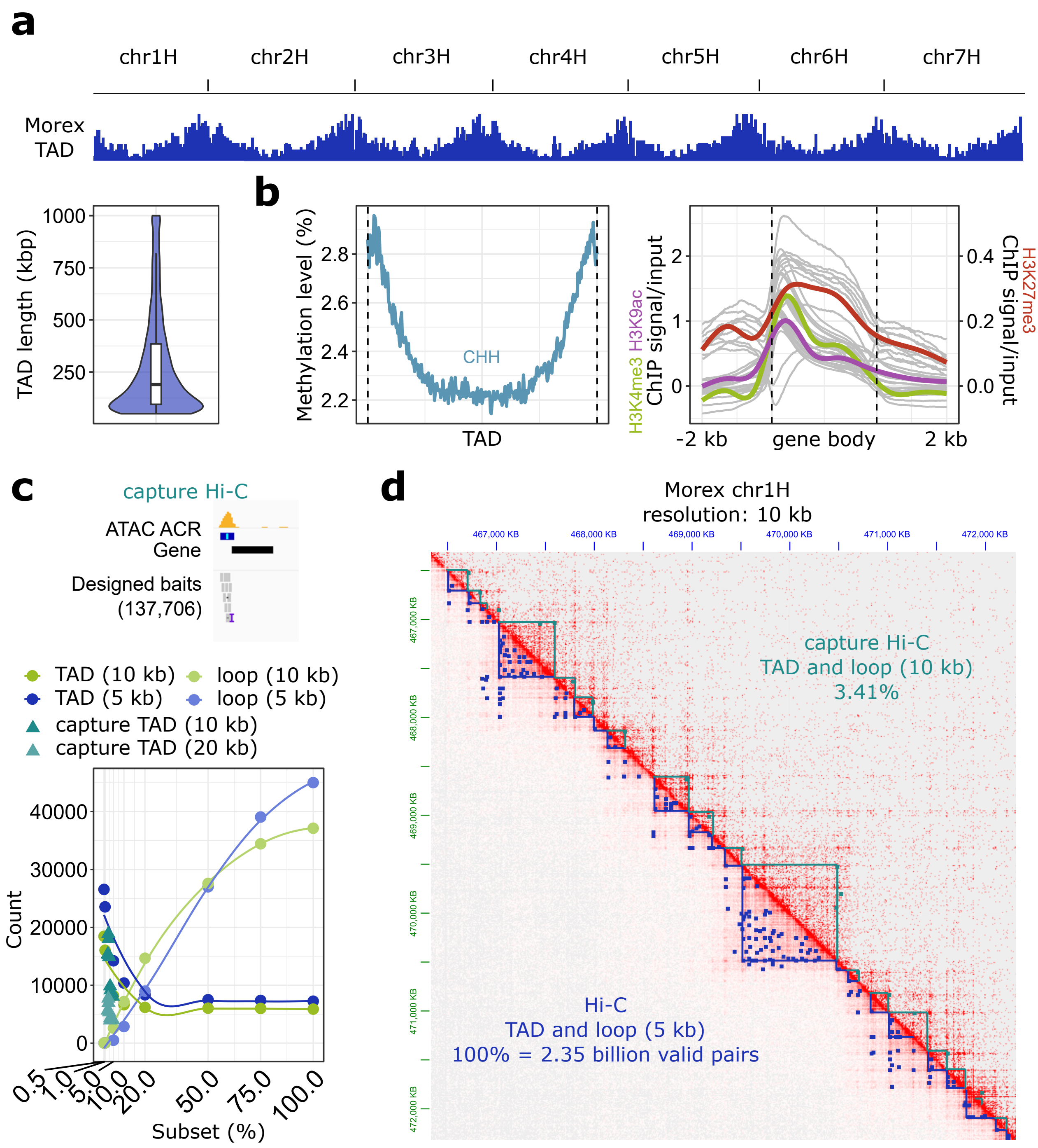


**Extended Data Fig. 5. Chromatin conformation assays identify barley TADs and loops.**
**a**, Chromosomal and length distribution of Morex TADs. Violin and boxplots show all TADs (n = 7,239). Boxes represent medians and interquartile ranges, and whiskers extend to 1.5x the interquartile ranges. **b**, Average CHH methylation profile of TADs in Morex. Average histone modification profiles across gene bodies; gray lines indicate individual genotypes (n = 10, per genotype average of biological replicates n = 2-3), and fitted curves show the overall trend. **c**, Capture Hi-C experiment based on 137,706 baits targeting the intersection of ACRs and 2 kb gene promoters. Impact of assay type, sequencing depth, and detection resolution on TAD and loop numbers. Fitted curves are based on in silico subsampling of Morex standard Hi-C data, whereas capture Hi-C data were from 10 genotypes (n = 10, **Supplementary Table 7**). **d**, Chromatin contact map of a representative region on Morex chr1H at 10 kb resolution, comparing TAD and loop annotations between capture Hi-C (upper panel) and standard Hi-C (lower panel).

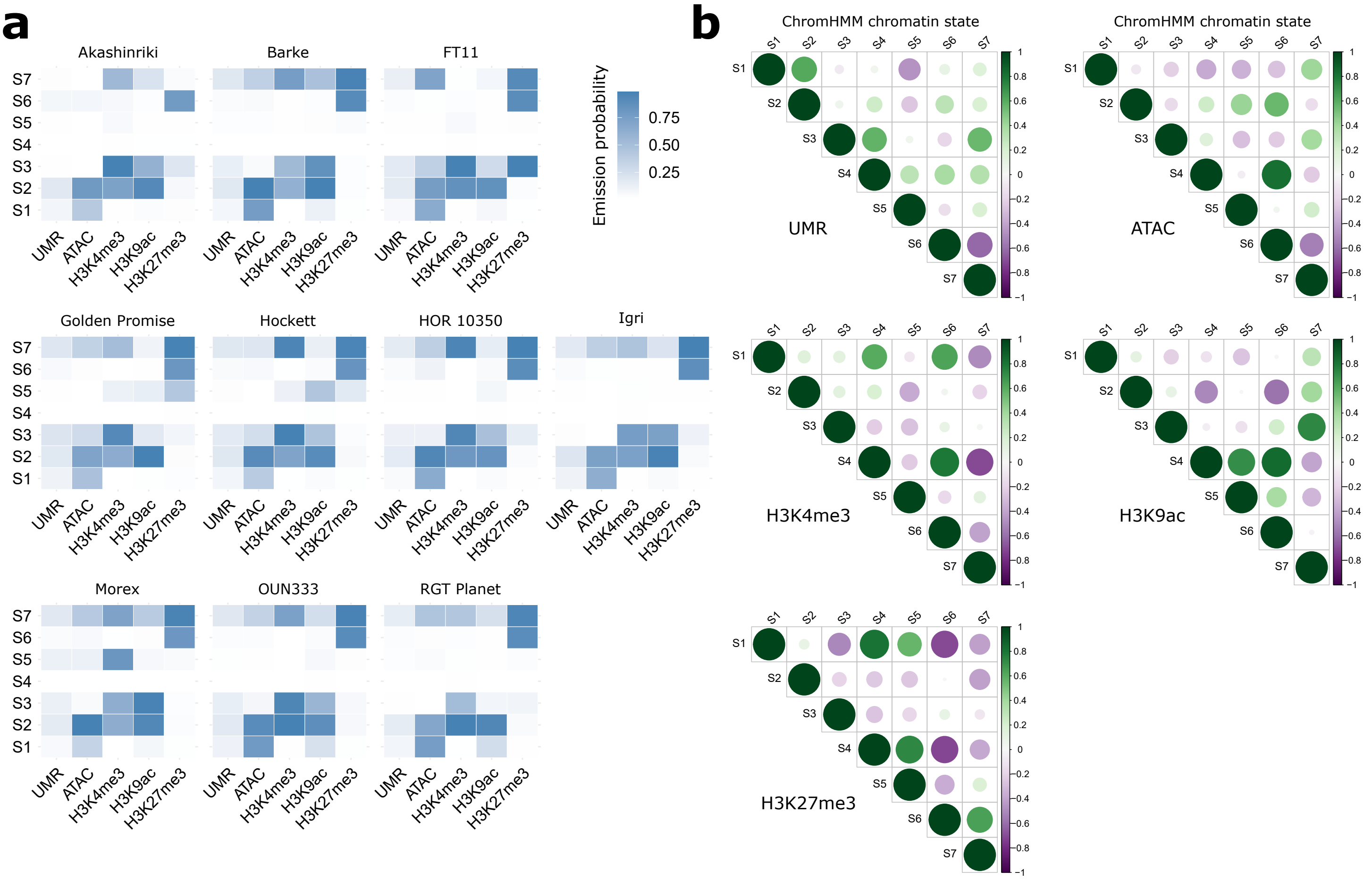


**Extended Data Fig. 6. ChromHMM-defined chromatin states and their conservation across 10 genotypes.**

**a**, Heatmaps show scaled (0-1) enrichment values of unmethylated regions (UMRs), ATAC-seq signal, H3K4me3, H3K9ac, and H3K27me3 across the seven chromatin states identified in 10 genotypes. **b**, Pairwise Pearson correlation coefficients of ChromHMM emission probabilities between seven chromatin states (S1-S7) across 10 genotypes for five epigenomic marks (UMRs, ATAC-seq signal, H3K4me3, H3K9ac, and H3K27me3. Higher coefficient values indicate greater conservation of chromatin state definitions across genotypes.

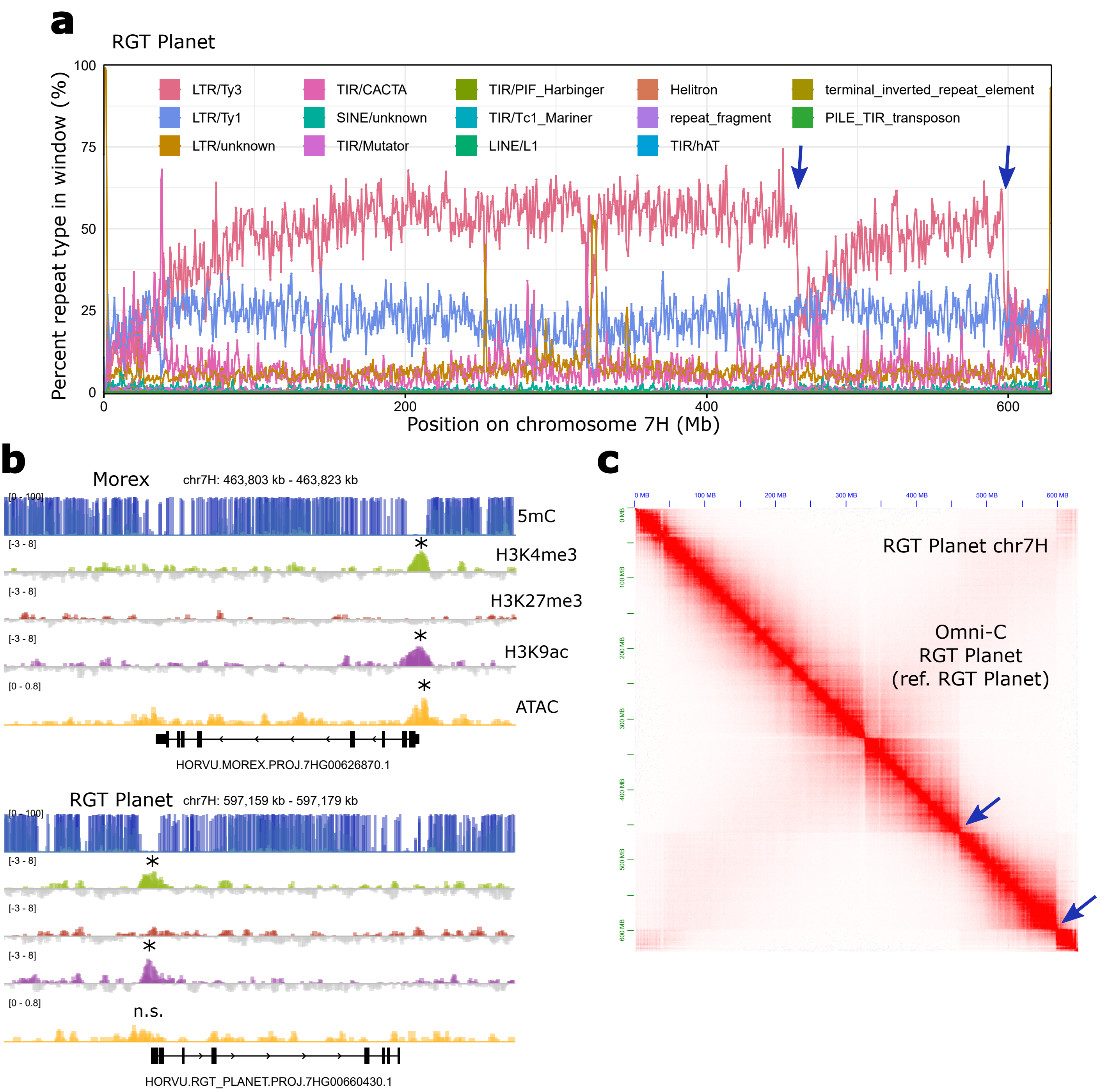


**Extended Data Fig. 7. Features of the chr7H inversion in RGT Planet.**
**a**, Distribution of TE types across RGT Planet chr7H (100 kb bins). **b**, Genome tracks of the first gene downstream of the inversion breakpoint in Morex and RGT Planet, showing DNA methylation (5mC, all contexts overlaid), histone modification signals, and chromatin accessibility; biological replicates are overlaid (ATAC-seq, n = 3; ChIP-seq, n = 2). Asterisks indicate significant peaks (q < 0.05). n.s., not significant. **c**, Omni-C map of chr7H in RGT Planet, mapped to the RGT Planet reference genome; blue arrows indicate the inversion breakpoints.

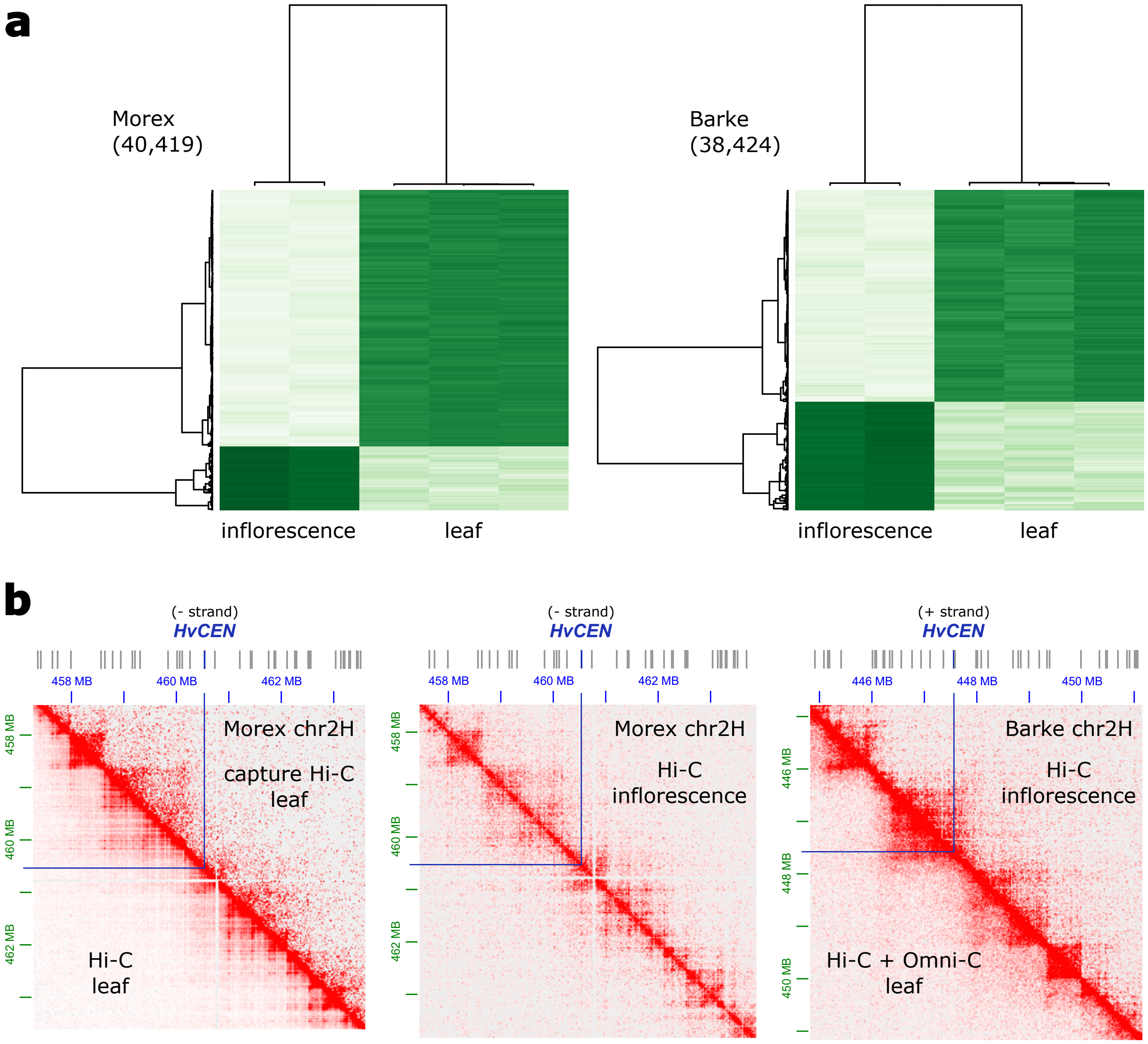


**Extended Data Fig. 8. Tissue-specific chromatin accessibility and chromatin interactions at the *HvCEN* locus.**
**a**, Heatmaps of tissue-specific accessible chromatin regions in Morex and Barke (FDR < 0.05), clustered by ATAC-seq signal, sample type, and biological replicate (leaf, n = 3; inflorescence, n = 2). Values and fold changes are provided in **Supplementary Data 7 and 8**. **b**, Contact maps of chr2H region in Morex and Barke (leaf and inflorescence) at 25 kb resolution, including the *HvCEN* locus (blue lines). All data were mapped to the genotype-specific reference genomes.

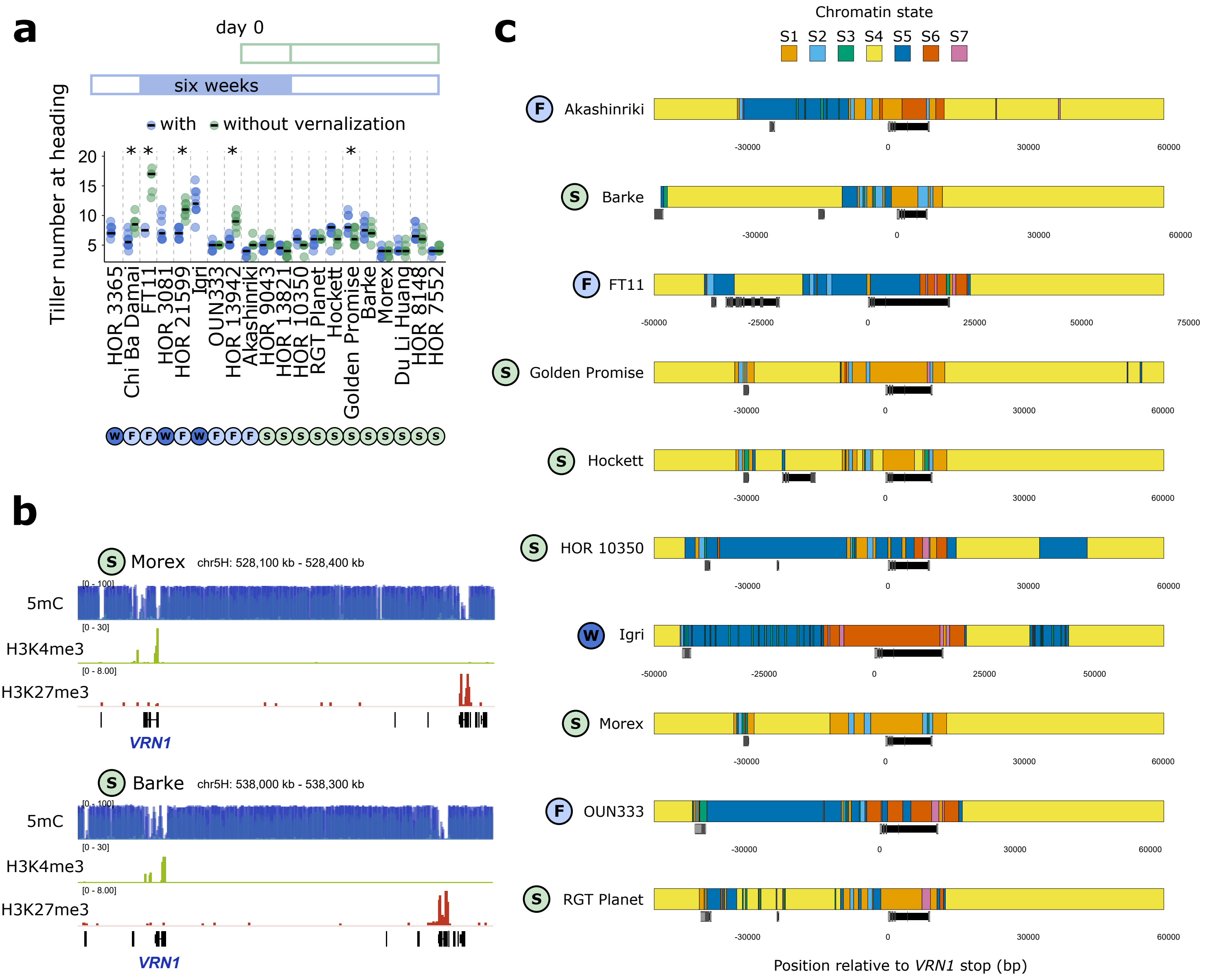


**Extended Data Fig. 9. Tiller number at heading and pre-vernalization chromatin states at the *VRN1* locus.**

**a**, Schematic of the flowering time experiment with and without six weeks of vernalization. The vernalization period was excluded when calculating days to heading. Total tiller number was recorded at heading. Horizontal bars indicate mean values (n = 2-10 biological replicates). Asterisks indicate significant differences between treatments ($P < 0.05$, two-way ANOVA, **Supplementary Table 9**). **b**, Genome tracks of a 300-kb window encompassing the *VRN1* locus in Morex and Barke, showing DNA methylation (5mC, all contexts overlaid), and histone modification signals (profiled by CUT&Tag). **c**, Pre-vernalization chromatin states at the *VRN1* locus across 10 genotypes. ChromHMM-derived chromatin state annotations (S1-S7) based on unmethylated regions (UMRs), ATAC-seq signal, H3K4me3, H3K9ac, and H3K27me3 across 10 genotypes. The winter barley Igri is characterized by an extensive repressive S6 chromatin state, whereas most spring barleys predominantly exhibit active S1/S2 states. Facultative genotypes have the silenced S6 segments interspersed with other chromatin states. Notably, the spring barley HOR 10350 shows facultative-like chromatin states despite being the only spring genotype among the ten that carries intact *VRN2*.

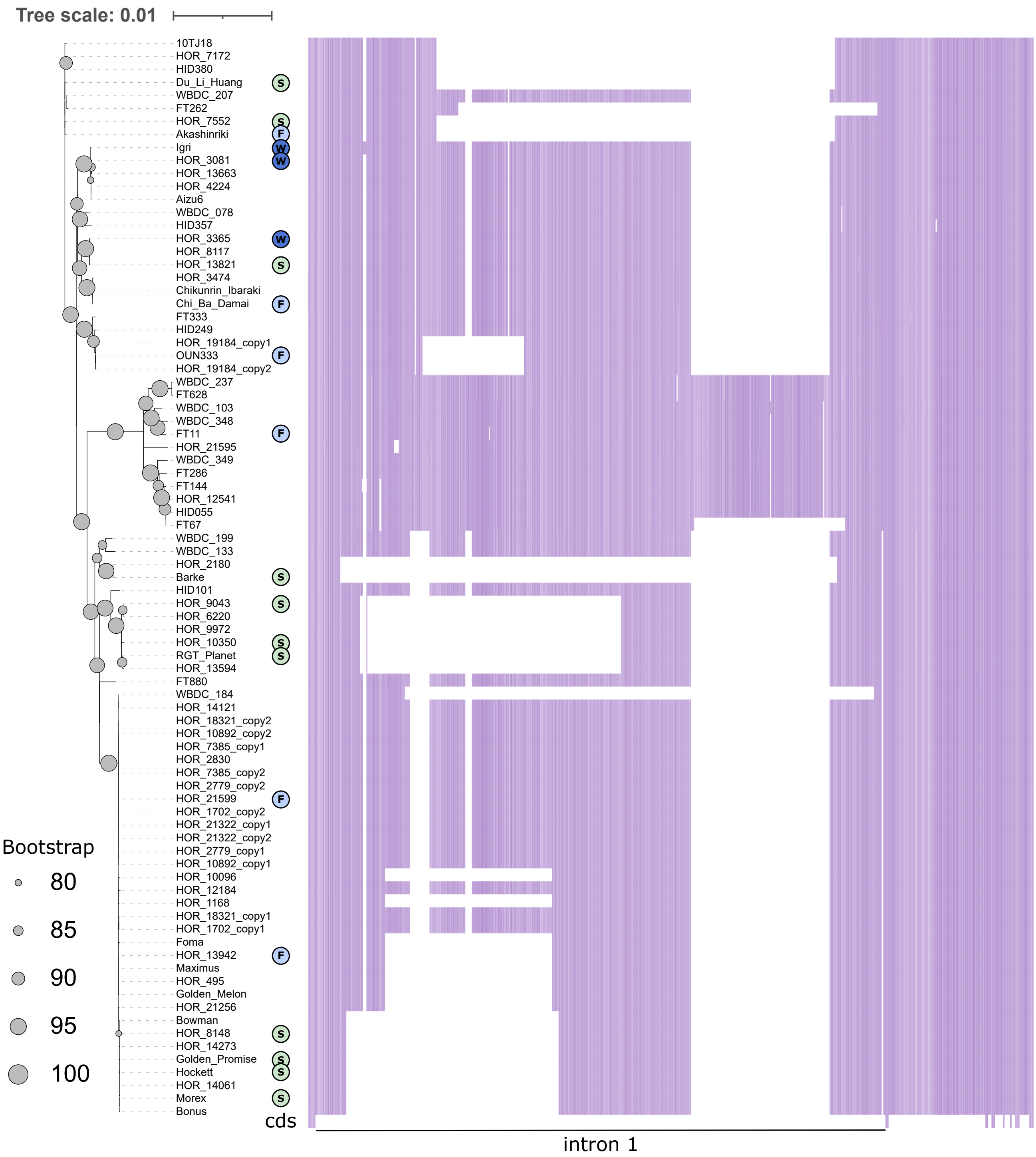


**Extended Data Fig. 10.** ***VRN1*** **allelic variation across 76 barley genotypes.**
Phylogenetic tree of *VRN1* genomic sequences across 76 barley genotypes, with sequence alignment to the Morex coding sequence; 20 genotypes have verified growth habit classifications **(Supplementary Table 10)**. The tree was generated with 1,000 bootstrap replicates (values ≥ 80 are shown).
